# Not all plant-based diets are alike: intake of minimally processed plant foods is associated with gut microbial richness among vegetarians and vegans

**DOI:** 10.64898/2026.11.19.767798

**Authors:** Andrés Méndez-Hurtado, Diego Coleto-Checa, Ane Alvarez, Helena Marcos-Pasero, Andrea Calderón, Alessandra Maestri, Alba Perez-Cuervo, Amaya Blanco-Rivero, Blanca Lacruz-Pleguezuelos, Nicolas Cárdenas-Roig, Lucía Carrasco-Guijarro, Adrián Martín-Segura, Elena Aguilar-Aguilar, Laura Judith Marcos-Zambrano

## Abstract

Plant-based diets are usually defined by what they exclude, overlooking which plant foods are eaten and how processed they are. Here we examine whether this overlooked dimension is related to the gut microbiome. In a cross-sectional study of 172 omnivorous, lacto-ovo-vegetarian and vegan adults profiled by shotgun metagenomics, vegetarians and vegans harboured fewer gut species than omnivores. Within this group, however, each standard deviation increase in minimally processed plant-food intake was associated with 27 additional species (p = 0.001), independently of fibre intake, and richness in participants with the highest intake approached that of omnivores. No such association was seen among omnivores. Higher intake was also linked to more species previously associated with favourable cardiometabolic health and to better health-related quality of life. Dietary labels alone may therefore conceal meaningful variation in the gut microbiome, and characterising how plant foods are processed may help in understanding how plant-based diets relate to health.

## 1. Introduction

Vegetarian and vegan diets have attracted growing scientific and public interest for their potential health benefits and lower environmental impact ^1–3^. These labels are not interchangeable: vegan diets exclude all animal-derived foods, whereas ‘vegetarian’ is an umbrella term encompassing lacto-ovo-, lacto-and ovo-vegetarian patterns, which differ in whether dairy, eggs or both are retained. Dietary recommendations in this direction, most prominently the Planetary Health Diet of the EAT–Lancet Commission, are explicit about food quality, placing whole grains, fruits, vegetables, legumes and nuts at the core of the diet ^2^. In practice, however, adopting a vegetarian or vegan diet only defines which foods are excluded, not which foods take their place: the same dietary label can encompass diets centred on minimally processed plant foods or relying heavily on refined and industrially formulated plant-derived products ^4,5^. In the NutriNet-Santé cohort, for example, vegans showed the highest ultra-processed food consumption of all dietary groups, close to 40% of energy intake, while at the same time scoring highest on established measures of plant-food quality ^4^. Plant-based diets should therefore not be considered nutritionally homogeneous.

This heterogeneity may be particularly relevant to the gut microbiome, a diet-responsive ecosystem involved in host metabolism ^6^. Plant foods supply fibre, microbiota-accessible carbohydrates and polyphenols that gut microbes ferment or transform ^7,8^, but the amount and accessibility of these substrates depend on the type of food and its degree of processing. If dietary labels group together diets that differ in these respects, comparisons between labels would be expected to yield inconsistent results, and this is largely what the literature shows. A systematic review of observational studies found no consistent association between vegan or vegetarian diets and gut microbial composition ^9^, and a review of intervention studies, while reporting some favourable shifts, found contradictory results for several taxa ^10^. Findings for alpha diversity are similarly discordant ^11^: vegetarians showed greater richness than omnivores in one amplicon-based study ^12^, no differences between vegans and omnivores were found in another using both amplicon and shotgun sequencing ^13^, and the largest metagenomic analysis to date, including 21,561 individuals, found lower richness in both vegans and vegetarians than in omnivores ^14^. In that study, associations between healthy plant-food intake and specific gut microbes were consistent across all three dietary patterns ^14^, suggesting that what is eaten within a pattern may matter as much as the pattern itself. Categorical comparisons may therefore be too coarse, as they cannot distinguish the effect of excluding animal products from that of the specific plant foods that replace them.

Plant-based diet indices are the most widely used tools for characterising plant-based eating beyond dietary labels. The overall plant-based diet index (PDI) captures the predominance of plant foods in the diet, while its healthful (hPDI) and unhealthful (uPDI) variants distinguish plant foods of higher and lower nutritional quality, and they show graded associations with type 2 diabetes and coronary heart disease ^15,16^. Their associations with gut microbial alpha diversity, however, have been inconsistent: hPDI has been positively associated with alpha diversity in some cohorts ^17^ but not in others, including a study using a modified hPDI ^18–20^, only when diet was assessed over the short term ^21^, or with evenness but not with species richness ^22^. All of these studies were conducted in predominantly omnivorous populations, so whether the indices capture variation in microbial diversity within vegetarian and vegan diets remains untested.

In mixed cohorts, these indices differ markedly between dietary patterns, with mean hPDI scores rising progressively from meat eaters to vegans ^4,14^, but how much variation they retain within each pattern has not been examined. Several features of their construction suggest that this within-pattern resolution may be limited. The indices were developed in predominantly omnivorous US cohorts, and food-group intakes are scored in quintiles of each study’s own distribution, an approach with recognised limitations ^23^; individuals who already avoid animal foods therefore tend to cluster in the upper score range, leaving less room to discriminate between them. More fundamentally, food groups were defined on the basis of nutrient and culinary similarity, and plant foods were classified as healthful or unhealthful according to prior evidence linking them to type 2 diabetes, cardiovascular disease, certain cancers and related conditions ^15^. Foods are therefore scored according to what they are made from rather than how they are made, which may explain how vegans can score highest on hPDI while also consuming the most ultra-processed foods ^4^. Many plant-based products now common in vegetarian and vegan diets, such as meat and dairy alternatives, also have no predefined place in the scoring framework, so their handling depends on study-specific decisions. This definition of quality was designed to predict disease risk, not microbial ecology, and other attributes along which plant foods vary, and which may be relevant to the gut microbiota, are not explicitly represented.

The degree of food processing is one such attribute. It is most widely operationalised through the NOVA classification, which assigns foods to four groups according to the extent and purpose of the industrial processing they undergo rather than their nutrient composition: unprocessed or minimally processed foods (group 1), processed culinary ingredients (group 2), processed foods (group 3), and ultra-processed foods (group 4), the latter defined as industrial formulations made predominantly from substances extracted from foods and from additives of exclusive industrial use ^24^. This axis is conceptually distinct from the nutritional-quality axis captured by plant-based diet indices, and the two can diverge in both directions. For example, olive oil and wholegrain bread are classified as healthful plant foods by hPDI yet fall outside NOVA group 1, whereas boiled potatoes, white rice and freshly squeezed fruit juice are minimally processed foods that the index scores as less healthful. Processing can disrupt the food matrix and chemically modify food components ^25^, and highly processed products may also contain ingredients and additives with potential effects on the gut microbiota ^26^. This evidence, however, largely concerns the ultra-processed end of the spectrum, and little is known about how variation in minimally processed plant-food intake relates to gut microbial diversity across dietary patterns.

To address this gap, we used data from the GutPlantHealth (GPH) study, a cross-sectional cohort of adults following omnivorous, lacto-ovo-vegetarian or vegan diets recruited in Madrid, Spain, in which gut microbiome and habitual diet were characterised using shotgun metagenomics and a food frequency questionnaire extended to capture plant-based foods and products, complemented by 3-day dietary records. We first compared microbial diversity across dietary patterns and examined whether established plant-based diet indices captured variation in microbial richness beyond dietary pattern. We then tested whether minimally processed plant-food (MPPF) intake, a continuous energy-adjusted measure, was associated with richness, whether this association depended on dietary pattern, and whether it was independent of fibre intake. We also characterised overall ultra-processed food intake across dietary patterns, and assessed microbial community composition and self-reported health-related quality of life as complementary outcomes. Finally, we examined whether the gut species associated with MPPF intake corresponded to species previously ranked as favourably or unfavourably associated with cardiometabolic health.

## 2. Materials and Methods

### 2.1 Study design and participants

This cross-sectional observational study (GutPlantHealth, GPH; protocol NUTRIGUT-MAD, IMD PI-069) was conducted at IMDEA Nutrition and Universidad Europea de Madrid (UEM) between March 2025 and January 2026. The study comprised an eligibility and enrolment visit (V0), followed by a single assessment visit (V1). Participants were recruited through social media channels associated with both institutions and through dissemination by professional dietetic associations and science communication networks.

Eligible participants were adults who self-reported following one of three dietary patterns for more than 12 months before enrolment: vegan, defined as excluding meat and all other animal-derived foods, including dairy products and eggs; lacto-ovo-vegetarian, defined as excluding meat while including both dairy products and eggs; or omnivorous, defined as consuming both plant-and animal-derived foods. Reported food intakes were reviewed and were broadly consistent with the assigned groups.

Recruitment prioritised balanced representation of the three dietary patterns. The enrolled population spanned 21–55 years, with body mass index ranging from 17.6 to 27.1 kg/m²; no participant had obesity. Age and body mass index were adjusted for in all subsequent models.

Exclusion criteria were psychiatric disease or an eating behaviour disorder; pregnancy or lactation; any disability or condition preventing adequate participation in the study procedures, or failure to meet the dietary patterns described; use of antibiotics, or of probiotic or prebiotic capsules, during the 3 months preceding enrolment; diagnosed gastrointestinal disease, such as Crohn’s disease, short bowel syndrome or irritable bowel syndrome; previous gastrointestinal surgery; and use of medications affecting the biochemical parameters analysed.

A total of 176 participants were enrolled (60 omnivores, 54 lacto-ovo-vegetarians and 62 vegans). Analytical sample sizes differ across analyses depending on data availability and on the exclusion criteria applied to each dietary exposure. These are specified in the corresponding sections and summarised in Figure 1. For the primary analyses, lacto-ovo-vegetarians and vegans were combined into a single plant-based group, as detailed in Section 2.10.1.

**Figure 1.**
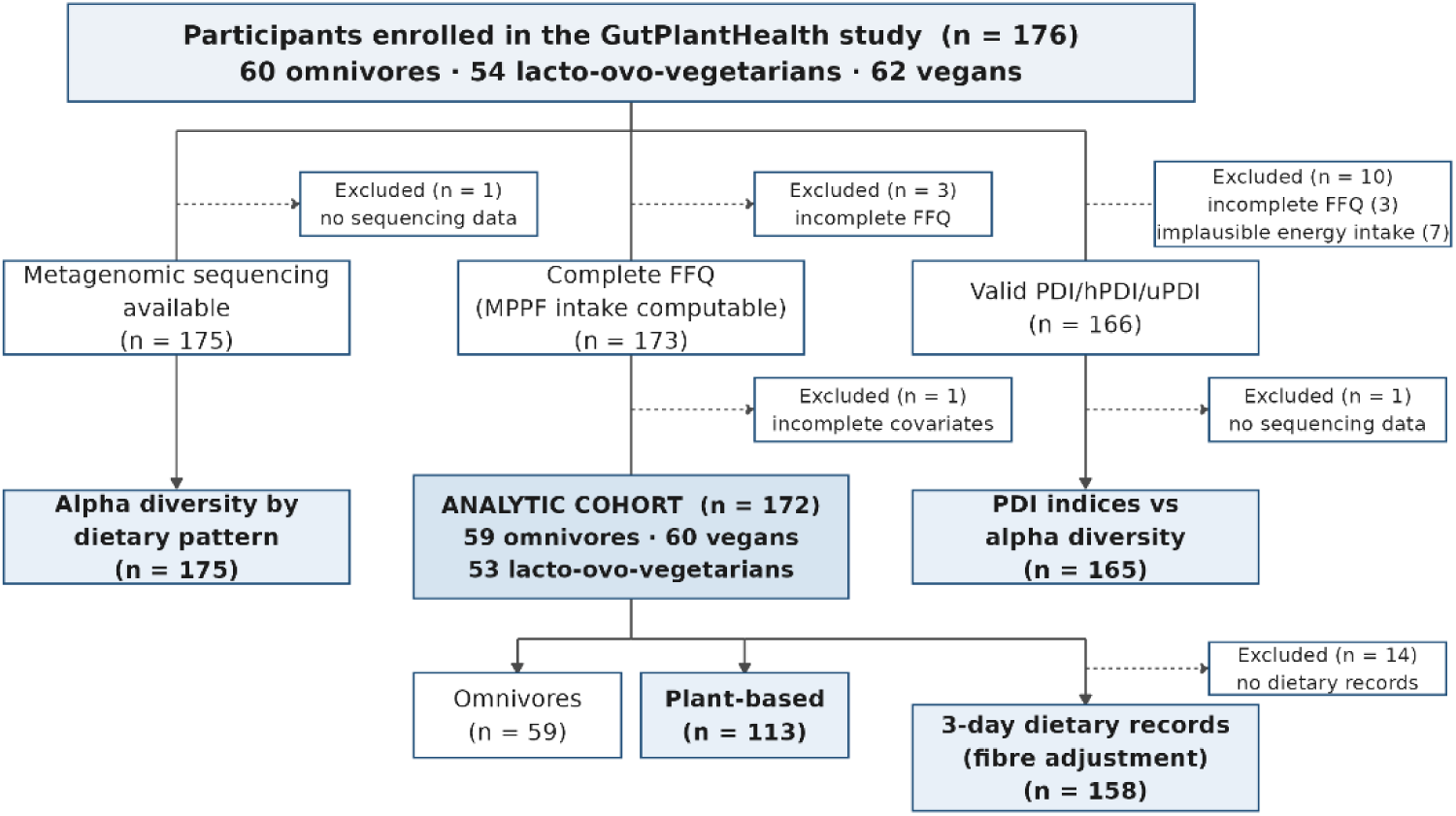
Participant flow through the study. Recruitment was by open call through social media and no formal eligibility-screening stage was recorded, so the diagram begins at enrolment. Analytic sample sizes differ across analyses because each requires different data: descriptive alpha-diversity comparisons require metagenomic data only (n = 175); MPPF intake models additionally require a complete food frequency questionnaire and complete covariate data (n = 172); and the plant-based diet indices additionally apply the energy-plausibility criterion of Satija et al. ^15^ (n = 166, of whom 165 had metagenomic data available). The analytic cohort (n = 172) was used for all MPPF intake analyses: species richness, overall microbial composition and self-reported quality of life. The subsample of the analytic cohort with 3-day dietary records (n = 158) was used to compare energy intake between the food frequency questionnaire and the records and to adjust for fibre intake. Dashed arrows indicate exclusions. FFQ, food frequency questionnaire; PDI, plant-based diet index; hPDI, healthful plant-based diet index; uPDI, unhealthful plant-based diet index.

### 2.2 Sample size

The sample size was determined a priori at the project design stage, based on the overall objective of detecting differences in gut microbial alpha diversity between dietary patterns. Given the absence of consistent estimates in previous comparisons of vegan, vegetarian and omnivorous populations ^9,27^, a medium effect size was assumed (f = 0.25) using Cohen’s conventional benchmark ^28^, an approach justified when no comparable estimates are available ^29^. Using the pwr package (version 1.3) ^30^ in R version 4.3.3 ^31^, for a one-way ANOVA across three groups with alpha diversity (Chao1 index) as the outcome, α = 0.05 and 80% power, 53 participants per group were required; allowing for an anticipated 15% drop-out rate, the target was set at 60 participants per group (180 in total).

### 2.3 Ethical considerations

The study was conducted in accordance with the Declaration of Helsinki and approved by the Research Ethics Committee of Fundación IMDEA Alimentación, currently Fundación IMDEA Nutrición (protocol “NUTRIGUT-MAD: Impacto de dietas basadas en plantas en el microbioma intestinal y la salud de la población de la Comunidad de Madrid”; approval code IMD PI-069, 14 January 2025). Signed informed consent was obtained from all participants at the enrolment visit (V0); this document included specific consent to microbial profiling. Data compiled during the study were pseudonymised in compliance with current Spanish and European legislation (Regulation (EU) 2016/679 and Organic Law 3/2018 on Personal Data Protection and Guarantee of Digital Rights). The biological sample collection was registered with the Spanish National Biobank Registry (Registro Nacional de Biobancos, Instituto de Salud Carlos III) under reference C.0008708.

### 2.4 Dietary assessment

Dietary intake was assessed in all participants using a semi-quantitative vegan-adapted food frequency questionnaire (vFFQ), designed to better capture plant-based diets while retaining the animal-derived food items of the original instrument, complemented by a 3-day dietary record ^32^. Dietary intake variables and plant-based diet indices were derived as described below.

#### 2.4.1 Food frequency questionnaire

Habitual dietary intake during the previous year was assessed using the vFFQ, an ad-hoc semi-quantitative questionnaire comprising 185 food and beverage items, self-administered electronically via REDCap ^33,34^ after standardised instructions provided by a registered dietitian. For each item, participants reported how often, on average, they had consumed the specified portion during the previous year, considering both consumption of the food alone and its inclusion in other foods or dishes. Responses were recorded using a nine-level frequency scale ranging from “never or <1 time/month” to “≥6 times/day”. Completed questionnaires were reviewed for missing or inconsistent responses, and participants were re-contacted when clarification was needed.

The questionnaire was adapted from the semi-quantitative FFQ validated in a Spanish population and used in the PREDIMED study ^35^, retaining its general structure and frequency-response scale. Of the 137 items in the original questionnaire, 125 were retained, and 12 were removed by collapsing redundant subcategories, mainly individual wine varieties and vegetable oils. Sixty new items were added to improve the representation of plant-based dietary patterns, resulting in a 185-item questionnaire. The new items covered plant-based dairy analogues (18 items), plant protein foods and meat alternatives (11), fermented foods (9), breads, cereals and tubers (8), nut and seed products differing in processing (6), beverages (4), and other items (4). The selection of additional items was informed by food categories represented in previous instruments and studies of vegan and plant-based populations ^36,37^. The vFFQ retains the portion definitions of the validated PREDIMED instrument; a validation study of the extended instrument is currently underway. The full item list is provided in Supplementary Table S1a.

Reported consumption frequencies were converted into servings per day. Servings were then converted into grams per day using predefined portion weights derived primarily from the original PREDIMED portion definitions and supplemented, when necessary, with Spanish portion-size reference tables ^38^. Energy intake was obtained by applying energy values per 100 g to the daily intake in grams of each item and summing across all items. Energy values were taken primarily from the Spanish Food Composition Database ^39^, supplemented by published Spanish food composition tables ^40^ and, for the few items without a suitable entry in these sources, manufacturer nutritional labels or international food composition databases. When a single vFFQ item represented more than one similar food, the assigned composition was the mean composition of the foods included in that item. Portion weights and energy values were reviewed and agreed upon by five registered dietitians. Item-level portion definitions and composition sources are provided in Supplementary Table S1b.

#### 2.4.2 Three-day dietary records

Participants additionally completed a 3-day dietary record at home, covering two weekdays and one weekend day, which was provided at one of the study visits. Records were entered by registered dietitians into the DIAL software (version 3.15; Alce Ingeniería, Madrid, Spain) ^41^, which estimates energy and nutrient intake from its own harmonised food composition table. Mean daily energy (kcal/day) and total dietary fibre (g/day) intakes over the 3 days were used, and fibre intake was additionally expressed per 1,000 kcal. Records were available for 158 of the 172 participants of the analytic cohort (Figure 1).

#### 2.4.3 Plant-based diet quality indices

The overall plant-based diet index (PDI), healthful plant-based diet index (hPDI), and unhealthful plant-based diet index (uPDI) were calculated following the scoring framework proposed by Satija et al. (2016, 2017). vFFQ items were assigned to the 18 food groups defined in the original methodology, with assignments reviewed by registered dietitians. Twenty-three vFFQ items were not included in the indices. Alcoholic beverages (n=3) and margarine (n=1) were excluded in accordance with the original methodology, which omits plant foods not consistently associated in one direction with health outcomes and margarine because of historical changes in its fatty acid composition ^15^. The remaining items had no counterpart among the 18 food groups: plant-based dairy analogues (n=7), other plant-derived products not represented in the original framework (n=2), condiments, spices and salt (n=6), and non-alcoholic or zero-calorie beverages (n=4). The complete item-to-group assignment is provided in Supplementary Table S1c.

Intake of each food group was adjusted for total energy intake using the residual method ^16,42^. A separate linear regression model was fitted for each of the 18 food groups, and the resulting residuals were added to the mean intake of the corresponding food group in the full cohort. Participants were then ranked into sex-specific quintiles of energy-adjusted intake, with cutoffs derived from the distribution of the full cohort, which included all three dietary pattern groups.

For PDI, plant food groups were assigned positive scores from 1 to 5 across increasing quintiles, whereas animal food groups were assigned reverse scores from 5 to 1. For hPDI, healthy plant food groups received positive scores, while less healthy plant and animal food groups received reverse scores. For uPDI, less healthy plant food groups received positive scores, while healthy plant and animal food groups received reverse scores. Scores across the 18 food groups were summed to obtain each index, with higher scores indicating greater adherence to the corresponding dietary pattern.

Energy intake was estimated using the food composition sources described in Section 2.4.1. In accordance with the criteria used by Satija et al. (2016), participants with implausible estimated energy intake were excluded from these analyses: <600 or >3,500 kcal/day for women and <800 or >4,200 kcal/day for men. Of the 176 enrolled participants, three were excluded because they did not complete the vFFQ and seven because of implausible energy intake, resulting in an analytic sample of 166 participants (Figure 1).

#### 2.4.4 Energy-adjusted intake of minimally processed plant foods (MPPF intake)

To complement the PDI, hPDI and uPDI with a measure focused on the processing level of plant foods, we derived a continuous measure of the energy-adjusted intake of minimally processed (NOVA group 1) plant foods, hereafter MPPF intake. Each of the 185 vFFQ items was classified individually according to the NOVA food-processing classification system ^24^ and assessed for vegan or plant-based status. Where an item’s processing level depended on commercial formulation rather than on the food itself, items were classified according to their predominant commercial presentation in the Spanish market. The complete item-level classification, including NOVA group, plant-based status, inclusion status and classification criterion, is provided in Supplementary Table S1d.

Vegan or plant-based items classified as NOVA group 1, corresponding to unprocessed or minimally processed foods, were included as MPPF items. These comprised 49 vFFQ items, including vegetables, fruits, legumes, whole grains, nuts and seeds, as well as unsweetened beverages such as coffee, tea and freshly squeezed juice. Animal-derived foods and plant-based items classified as NOVA groups 2–4 were excluded.

The primary measure applied this commercial criterion throughout and therefore excluded five items whose processing level depends on commercial formulation rather than on the food itself: tofu, tempeh and natto (typically marketed with added coagulants, salt or seasonings), roasted peanuts (typically salted) and textured soy protein. In a sensitivity analysis these five items were treated as minimally processed, as they would be in artisanal or additive-free preparations, yielding a broader 54-item version of the measure.

For each participant, intake of each included item was expressed as grams per 1,000 kcal by dividing daily intake in grams by total daily energy intake and multiplying by 1,000. MPPF intake was calculated by summing the energy-adjusted intake of the 49 included items. The resulting value was standardised within the analytic sample as a z-score for use in statistical models, such that a one-unit increase represented one standard deviation. Three participants who did not complete the vFFQ were excluded (Figure 1).

#### 2.4.5 Ultra-processed food intake

To characterise the overall processing level of the diet independently of its plant-based content, we calculated ultra-processed food (UPF) intake as the proportion of daily energy derived from vFFQ items classified as NOVA group 4, using the item-level classification and composition sources described above (Supplementary Tables S1b and S1d). Energy from NOVA group 4 items was divided by total daily energy intake and expressed as a percentage, the standard metric in the ultra-processed food literature. This variable was computed identically in all participants and does not depend on whether an item is of plant or animal origin.

### 2.5 Health-related quality of life

Health-related quality of life was assessed at the evaluation visit (V1) using the Spanish-language 36-Item Short Form Health Survey (SF-36), self-administered electronically through REDCap. Item responses were recoded and transformed to 0–100 scales following the standard SF-36 scoring algorithm ^43,44^, yielding eight dimension scores: general health, physical functioning, role–physical, role– emotional, social functioning, bodily pain, vitality, and mental health, with higher scores indicating better health-related quality of life. Dimension scores were computed following the RAND missing-data rule and combined without imputation. An overall SF-36 score was calculated as the unweighted mean of the eight dimensions and was available for all 172 participants in the analytic cohort.

### 2.6 Anthropometric and biochemical assessment

Height was measured using a stadiometer (TANITA HR-001), while body weight and body composition, including body fat percentage and visceral fat rating, were assessed by bioelectrical impedance analysis using a TANITA RD-545 device, following the manufacturer’s standardised procedures. Body mass index (BMI) was calculated as body weight in kilograms divided by height in metres squared.

At the evaluation visit (V1), venous blood samples were collected after an overnight fast and were measured according to a standardised protocol by Laboratory CQS Consulting (UNE-ISO 15189:2007, No. 659/LE1318). The biochemical assessment included haemoglobin, vitamin B12, folate, and vitamin D as markers related to nutritional status, together with glucose, glycated haemoglobin (HbA1c), total cholesterol, triglycerides, high-density lipoprotein cholesterol, and low-density lipoprotein cholesterol as markers of glycaemic and lipid status. Of the 41 biochemical parameters measured, 22 are reported in Table 1 and the remaining 19 in Supplementary Table S2.

**Table 1.** Baseline characteristics, dietary intake and biochemical profile by dietary pattern. Values are mean (SD) for continuous variables, median [IQR] for insulin, HOMA-IR, CRP, triglycerides and ultra-processed food intake, and n (%) for categorical variables. Dietary intake was estimated from the food frequency questionnaire; ultra-processed foods are NOVA group 4 items; PDI, hPDI and uPDI are available for 165 participants. p-values from Kruskal–Wallis tests for continuous variables and chi-squared tests for categorical variables; q-values are Benjamini–Hochberg corrected across the 41 biochemical parameters analysed. n.a., not applicable; HOMA-IR, homeostatic model assessment of insulin resistance; CRP, C-reactive protein; PDI, plant-based diet index; hPDI, healthful PDI; uPDI, unhealthful PDI.

| Characteristic | Omnivores (n=59) | Lacto-ovo-vegetarians (n=53) | Vegans (n=60) | p | q |
|---|---|---|---|---|---|
| **Demographics and anthropometry** |  |  |  |  |  |
| Age, years | 30.8 (6.7) | 32.3 (7.7) | 36.6 (8.4) | <0.001 | n.a. |
| Female sex | 29 (49%) | 16 (30%) | 24 (40%) | 0.124 | n.a. |
| Current smoker | 14 (24%) | 14 (26%) | 17 (28%) | 0.848 | n.a. |
| Weight, kg | 66.1 (10.5) | 62.0 (9.6) | 63.6 (9.6) | 0.135 | n.a. |
| BMI, kg/m<sup>2</sup> | 22.7 (2.1) | 22.2 (2.2) | 22.2 (2.2) | 0.293 | n.a. |
| Body fat, % | 22.6 (6.1) | 24.4 (6.3) | 23.8 (7.1) | 0.253 | n.a. |
| Visceral fat rating | 3.9 (2.3) | 3.6 (2.4) | 4.3 (2.5) | 0.272 | n.a. |
| **Dietary intake** |  |  |  |  |  |
| Energy intake, kcal/day | 2352 (1019) | 2153 (636) | 2531 (856) | 0.040 | n.a. |
| MPPF intake, g/1,000 kcal | 580 (207) | 689 (222) | 645 (188) | 0.038 | n.a. |
| Ultra-processed foods, % of energy | 19.7 [12.5–25.0] | 20.8 [14.6–27.1] | 22.1 [16.7–31.7] | 0.061 | n.a. |
| PDI | 45.5 (5.7) | 54.7 (5.0) | 61.1 (4.5) | <0.001 | n.a. |
| hPDI | 46.2 (7.8) | 52.9 (8.7) | 62.8 (7.8) | <0.001 | n.a. |
| uPDI | 50.4 (6.4) | 55.3 (6.9) | 57.1 (6.6) | <0.001 | n.a. |
| **Nutritional status** |  |  |  |  |  |
| Haemoglobin, g/dL | 14.4 (1.1) | 14.0 (1.0) | 13.8 (1.5) | 0.049 | 0.118 |
| Haematocrit, % | 43.7 (3.2) | 42.3 (3.0) | 41.8 (4.0) | 0.012 | 0.032 |
| MCV, fL | 90.5 (4.2) | 90.8 (4.7) | 90.3 (6.4) | 0.857 | 0.878 |
| RDW, % | 13.6 (0.6) | 13.6 (1.0) | 13.8 (1.3) | 0.701 | 0.822 |
| Vitamin B12, pg/mL | 433.4 (164.4) | 443.8 (177.9) | 552.9 (224.2) | 0.003 | 0.013 |
| B12 supplementation | 0 (0%) | 35 (66%) | 53 (88%) | <0.001 | n.a. |
| Folate, ng/mL | 8.0 (2.6) | 8.7 (2.8) | 10.5 (3.3) | <0.001 | <0.001 |
| Vitamin D (25-OH), ng/mL | 23.3 (8.2) | 26.9 (8.9) | 26.2 (14.4) | 0.161 | 0.291 |
| Albumin, g/dL | 4.59 (0.29) | 4.46 (0.24) | 4.41 (0.30) | 0.004 | 0.013 |
| Prealbumin, mg/dL | 28.1 (4.0) | 25.3 (3.2) | 24.4 (4.2) | <0.001 | <0.001 |
| Calcium, mg/dL | 9.8 (0.3) | 9.7 (0.3) | 9.6 (0.3) | <0.001 | 0.004 |
| **Glycaemic profile** |  |  |  |  |  |
| Glucose, mg/dL | 70.9 (6.2) | 72.0 (7.0) | 71.2 (8.9) | 0.625 | 0.822 |
| HbA1c, % | 5.2 (0.2) | 5.2 (0.2) | 5.2 (0.3) | 0.820 | 0.868 |
| Insulin, µIU/mL | 3.60 [2.75–4.90] | 3.60 [2.70–4.50] | 3.40 [2.68–4.43] | 0.826 | 0.868 |
| HOMA-IR | 0.60 [0.50–0.90] | 0.60 [0.50–0.80] | 0.60 [0.50–0.80] | 0.647 | 0.822 |
| **Lipid profile** |  |  |  |  |  |
| Total cholesterol, mg/dL | 184.1 (33.7) | 161.2 (33.1) | 151.8 (24.1) | <0.001 | <0.001 |
| LDL cholesterol, mg/dL | 106.8 (28.2) | 84.8 (27.8) | 79.2 (21.2) | <0.001 | <0.001 |
| HDL cholesterol, mg/dL | 64.0 (13.8) | 62.7 (14.0) | 59.1 (11.4) | 0.163 | 0.291 |
| Triglycerides, mg/dL | 62 [44–81] | 63 [50–80] | 61 [52–75] | 0.782 | 0.867 |
| **Inflammation and immune cells** |  |  |  |  |  |
| C-reactive protein, mg/dL | 0.10 [0.00–0.10] | 0.10 [0.00–0.10] | 0.10 [0.00–0.10] | 0.940 | 0.940 |
| TNF-α, pg/mL | 7.0 (3.0) | 6.8 (2.8) | 7.3 (2.9) | 0.764 | 0.867 |
| Leukocytes, ×10<sup>3</sup>/µL | 6.1 (1.6) | 5.5 (1.4) | 4.9 (1.5) | <0.001 | <0.001 |
| Lymphocytes, cells/µL | 2151 (473) | 1788 (461) | 1613 (448) | <0.001 | <0.001 |

### 2.7 Faecal sample collection, DNA extraction, and sequencing

Faecal samples were self-collected by participants using sterile collection tubes containing a DNA stabilisation buffer and an integrated collection spoon (Canvax Reagents, S.L.U.; catalogue no. SC0010), following written instructions provided at enrolment. Samples were transported to the laboratory by participants under refrigerated conditions using ice packs and were stored at −80 °C until DNA extraction.

Total DNA was extracted from faecal samples using the QIAamp Fast DNA Stool Mini Kit (QIAGEN; catalogue no. 51604), following the manufacturer’s protocol. Shotgun metagenomic sequencing was performed by Novogene Europe (Munich, Germany) on the Illumina NovaSeq X Plus platform using 150-bp paired-end sequencing, targeting approximately 6 Gb of raw sequence data per sample. Samples were sequenced in two batches (n = 96 and n = 79). Raw FASTQ files were provided by the sequencing provider for downstream preprocessing and analysis.

### 2.8 Metagenomic data processing and taxonomic profiling

#### Read preprocessing

Raw paired-end reads were quality-trimmed using fastp (version 0.25.0) ^45^, including front-end trimming, a minimum retained read length of 50 bp, and a quality threshold of Q20. Human-derived reads were subsequently removed by alignment against the human reference genome GRCh38 using Bowtie2 (version 2.5.4) ^46^, and host-depleted read pairs were retained for downstream analysis.

#### Sequencing depth

Read counts and quality statistics were computed with SeqKit (version 2.10.0) ^47^. One sample showed markedly greater sequencing depth than the remainder of the cohort (594 million reads, ∼89 Gb of raw sequence data) and was randomly downsampled to 28.0 million read pairs using seqkit sample, yielding a depth representative of the remaining samples. Across the final set of 175 samples, mean sequencing depth was 28.8 ± 13.4 million read pairs (median 25.5).

#### Taxonomic profiling

Host-depleted reads were taxonomically profiled using MetaPhlAn 4 (version 4.2.2) with the mpa_vJan25_CHOCOPhlAnSGB_202503 database ^48^. Per-sample profiles were merged using the MetaPhlAn merge_metaphlan_tables.py utility and converted from species-level genome bin taxonomy to Genome Taxonomy Database taxonomy, release 220 ^49^, using the sgb_to_gtdb_profile.py conversion script. Polyphyletic lineages were represented using standard GTDB alphabetical suffixes, such as *Bacillota_A* to *Bacillota_I*. The resulting GTDB-resolved taxonomic profile was used for all diversity and community-composition analyses.

### 2.9 Data processing and quality control

#### Data import and object construction

GTDB-resolved taxonomic profiles were imported into R as a TreeSummarizedExperiment object using the mia package (version 1.10.0) ^50^. Taxonomic annotations from domain to species level were parsed from each taxon’s lineage. Analyses were restricted to taxa with an informative species-level annotation, yielding 2,254 species.

#### Screening for potential batch-associated contaminants

Given that samples were sequenced in two batches (n = 96 and n = 79), all species-level taxa were screened for marked differences in prevalence between batches. Taxa showing ≥80% prevalence in one batch and ≤20% prevalence in the other were flagged as potential batch-associated artefacts. Two species met this criterion, both showing complete batch specificity (100% versus 0% prevalence).

To identify additional taxa associated with sequencing batch, prevalence was compared between batches using Fisher’s exact tests, with Benjamini–Hochberg correction for multiple testing. Twelve species were significantly associated with sequencing batch at q < 0.05, including the two species identified by the prevalence-based screening. Each of these taxa was individually evaluated based on published ecological and taxonomic evidence.

Five species were excluded as potential contaminants because they showed strong batch association and were considered implausible members of the human gut microbiota: *Microbacterium ginsengisoli* and *Microbacterium* sp001619615, belonging to a genus previously reported as a common reagent or laboratory contaminant ^51^; *Sphingomonas* sp017418975 and *Sphingomonas* sp005503355, also belonging to a contaminant-associated genus ^51^; and *Sediminibacterium* sp902168225, a freshwater-associated organism with no documented evidence of human gut colonisation ^52,53^. The remaining seven batch-associated taxa were retained because they represented documented or plausible gut-associated lineages or lacked sufficient ecological evidence to support exclusion. After filtering, 2,249 species were retained for all downstream diversity and community-composition analyses.

### 2.10 Statistical analysis

#### 2.10.1 Alpha diversity

Alpha diversity, including observed species richness, Shannon diversity, and Pielou’s evenness, was calculated at the species level using the mia package (version 1.10.0) ^50^ on the 2,249 species retained after quality-control filtering.

##### Dietary-pattern comparisons

Differences in alpha-diversity metrics among the three dietary-pattern groups were assessed in all participants with available metagenomic data (n=175), using Kruskal–Wallis tests followed by Dunn’s pairwise tests with Benjamini–Hochberg correction.

##### Plant-based diet indices

Because PDI, hPDI, and uPDI are cohort-relative indices constructed using quintile-based scoring, their associations with observed species richness were assessed using partial Spearman correlations (ppcor package, version 1.1; ^54^) adjusted for sex, age, BMI, and sequencing depth, both in the pooled sample and within each dietary pattern, and with additional adjustment for dietary pattern. Benjamini–Hochberg correction was applied separately to pooled and stratified analyses.

As a complementary measure, a continuous healthful plant-food intake variable (hPDI_raw) was calculated by summing energy-adjusted intake (g/1,000 kcal) across the healthful plant-food categories defined by Satija et al. ^15,16^.

##### Between- and within-pattern variation

To assess whether each dietary measure varied mainly between or within dietary patterns, the proportion of the variance of PDI, hPDI, uPDI, hPDI_raw and MPPF intake explained by dietary pattern (η²) was estimated from one-way analysis of variance, and within-pattern variability was described using the coefficient of variation in each dietary pattern.

##### MPPF intake

The primary analysis compared omnivores with plant-based participants, combining lacto-ovo-vegetarians and vegans into a single group (n=113), as both patterns exclude meat and were hypothesised to share the exposure of interest. The association between MPPF intake and observed species richness was assessed using linear regression models including a dietary pattern × MPPF intake interaction term, and models stratified by dietary pattern, adjusting for sex, age, BMI, and sequencing depth. Models stratified by the three original dietary patterns are reported as secondary analyses.

##### Exploratory categorical analysis

As an exploratory analysis, observed species richness was compared among omnivores and plant-based participants categorised into tertiles of MPPF intake. Differences among the four groups were assessed using Kruskal–Wallis tests followed by Dunn’s pairwise tests with Benjamini–Hochberg correction.

##### Sensitivity analyses

Robustness of the MPPF intake–richness association was assessed within plant-based participants, within omnivores and for the dietary pattern × MPPF intake interaction, using: (i) a broader MPPF definition additionally including the five items whose processing level depends on commercial formulation, resulting in a 54-item version as described in Section 2.4.4; (ii) re-analysis with richness recomputed after excluding species detected below relative-abundance thresholds of 0.001%, 0.01%, 0.1% and 1%, applied within each sample; (iii) robust regression using Huber M-estimation, implemented with rlm from the MASS package (version 7.3-60.0.1) ^55^, with p values approximated from t statistics, and MM-estimation with lmrob from the robustbase package (version 0.99-4-1) ^56^ as a check; (iv) leave-one-out analyses omitting, in turn, each of the 49 MPPF items and each participant; and (v) additional adjustment for total fibre intake (g/1,000 kcal) estimated from the 3-day dietary records, in the subsample with records available, with models without fibre refitted in the same subsample for comparison. To assess whether results depended on quantifying intake by weight, MPPF intake was also expressed as servings/1,000 kcal, converting the FFQ frequency categories of the same 49 items to servings per day.

##### Ultra-processed food intake

Differences in ultra-processed food intake across dietary patterns were assessed using Kruskal–Wallis tests, and the comparison between omnivores and plant-based participants was additionally modelled by linear regression adjusting for sex, age and BMI. The association between MPPF intake and ultra-processed food intake was assessed using Spearman correlations within each dietary pattern.

##### Comparison with dietary records

Agreement between energy intake estimated by the vFFQ and by the 3-day dietary records was assessed using Spearman correlation, the mean difference between methods and cross-classification into quartiles.

##### Multiplicity

A single primary analysis was defined for each outcome (omnivores versus plant-based participants). All remaining models are reported with nominal p-values and interpreted as hypothesis-generating.

#### 2.10.2 Microbial community composition

Prior to community-composition analyses, species were filtered to those present at a relative abundance above 0.01% in at least 10% of samples, a prevalence threshold shown to yield more consistent results in benchmarking of microbiome analysis methods ^57^, yielding 408 species. The abundance threshold was added to limit the influence of trace-level detections. This filter was applied only to community-composition analyses; alpha-diversity metrics were computed on unfiltered profiles.

Community composition was assessed using Aitchison distance, calculated as Euclidean distance on centred log-ratio-transformed species-level relative abundances. Centred log-ratio transformation was performed using transformAssay from the mia package, after replacing zeros with a pseudocount corresponding to half the minimum non-zero relative-abundance value observed across the dataset.

Differences in overall community composition among dietary-pattern groups were assessed using permutational multivariate analysis of variance (PERMANOVA), implemented with adonis2 from the vegan package (version 2.7-1) ^58^ using marginal tests, with models adjusted for MPPF intake, sex, age and BMI. Pairwise comparisons between dietary patterns were performed with the same model, with Benjamini–Hochberg correction. Homogeneity of multivariate dispersion among dietary patterns was assessed using PERMDISP (betadisper, vegan package) followed by Tukey’s honest significant difference test.

Associations between MPPF intake and community composition were assessed using PERMANOVA models adjusted for dietary pattern, sex, age and BMI in the full cohort, and fitted separately within plant-based participants and within omnivores with adjustment for sex, age and BMI. Aitchison distance was the primary metric. As a sensitivity analysis to the treatment of zero values, analyses were repeated using robust centred log-ratio transformation (rCLR) ^59^, which calculates geometric means using non-zero components without adding a pseudocount. Additional analyses using Jaccard and Bray–Curtis dissimilarities were conducted to examine whether associations were more closely related to taxon presence or absence or to abundance structure. In the stratified analyses, Benjamini–Hochberg correction was applied across the eight tests (two strata × four metrics). All PERMANOVA and PERMDISP analyses used 9,999 permutations.

#### 2.10.3 Health- and diet-associated gut species

To examine whether the species associated with MPPF intake corresponded to species previously associated with host health, we used the ZOE Microbiome Health Ranking 2025 ^60^, which ranks 661 gut species according to their associations with cardiometabolic health markers and with diet quality in more than 34,000 individuals. For each ranking, the 50 most favourably and the 50 most unfavourably ranked species were defined a priori as species sets; the 100 most favourably and unfavourably ranked species were used in a sensitivity analysis. Because the rankings are defined at the species-level genome bin (SGB) level, SGB-level profiles from the same MetaPhlAn run described in Section 2.8 were used for this analysis, and SGBs merged by MetaPhlAn into SGB groups were matched by their SGB identifier.

For each participant, the number of species of each set present (relative abundance > 0) was modelled as a function of MPPF intake using quasi-Poisson regression adjusted for sex, age, BMI and sequencing depth, fitted separately within plant-based participants and omnivores. To assess whether these associations exceeded those expected from the overall increase in species richness, the association of each set with MPPF intake was compared with that of 1,000 random sets of species with similar prevalence. Statistical significance of this difference was assessed by permutation of MPPF intake across participants (9,999 permutations, preserving the associations with covariates), an approach that accounts for co-occurrence among species. Robustness of the association with health-favourable species was assessed using species-presence thresholds of 0.001% and 0.01% relative abundance, additional adjustment for vegan versus lacto-ovo-vegetarian pattern, and leave-one-participant-out analyses.

As an exploratory analysis, the presence of each species of the health sets with a prevalence between 10% and 90% within the stratum was modelled using logistic regression with the same covariates, with Benjamini-Hochberg correction within each set and stratum. Species were named according to GTDB taxonomy (release 220, as in Section 2.8) using the MetaPhlAn SGB-to-GTDB mapping.

#### 2.10.4 Health-related quality of life and biochemical parameters

The association between MPPF intake and the overall SF-36 score was assessed using linear regression models including a dietary pattern × MPPF intake interaction term and adjusting for sex, age, and BMI. As in Section 2.10.1, the primary contrast compared omnivores with the combined plant-based group, and models stratified by the three original dietary patterns were fitted as secondary analyses. The correlation between species richness and the overall SF-36 score was assessed using Spearman correlation. Associations of MPPF intake and of PDI, hPDI and uPDI with biochemical and anthropometric parameters were assessed using partial Spearman correlations adjusted for sex, age and BMI, with and without additional adjustment for dietary pattern and within each dietary pattern, with Benjamini-Hochberg correction.

### 2.11 Use of large language models

Claude Opus 5.5 (Anthropic) was used to assist in writing and debugging the scripts for data processing, statistical analysis and figure generation, and in drafting and editing the manuscript text. All figures are plots produced by this code directly from the study data; no generative AI images were used. The authors designed the study and the analyses, checked the code and its outputs, verified all reported results against the source data and all references against the original publications, and revised the text. The authors take full responsibility for the content of the manuscript, the analyses and the code (see Code availability).

## 3. Results

### 3.1 Dietary patterns are associated with distinct biochemical profiles despite similar anthropometric characteristics

Of 176 enrolled participants, 172 (59 omnivores, 53 lacto-ovo-vegetarians and 60 vegans; 113 plant-based) comprised the analytic cohort (Figure 1). Baseline characteristics, dietary intake and biochemical profile by dietary pattern are shown in Table 1. Age differed across dietary patterns (p<0.001), with vegans older than omnivores (36.6 vs 30.8 years), a difference adjusted for in all subsequent models; anthropometric and body-composition measures did not differ across groups. Sixteen of 41 biochemical parameters differed across dietary patterns after Benjamini-Hochberg correction (Table 1 and Supplementary Table S2). Lipid parameters were progressively lower from omnivores to vegans (total cholesterol 184.1 vs 151.8 mg/dL, LDL cholesterol 106.8 vs 79.2 mg/dL, and the total cholesterol/HDL and LDL/HDL ratios; all q<0.05). Vitamin B12 and folate were higher in vegans, the former reflecting a substantially higher prevalence of B12 supplementation, whereas albumin, prealbumin and calcium were modestly lower. Red-cell indices showed small reductions in haematocrit and red blood cell count, and all leukocyte subpopulations (total leukocytes, lymphocytes, monocytes and neutrophils) were lower in vegans. All reported values remained within clinical reference ranges. Among the 158 participants with 3-day dietary records, energy intake estimated by the vFFQ correlated with that estimated from the records (Spearman ρ = 0.37, p < 0.001), with a mean difference of 233 kcal/day; 38% of participants were classified into the same quartile of energy intake by both methods and only 5% into the opposite quartile.

### 3.2 Gut microbial richness is lower in plant-based dietary patterns, and plant-based diet indices largely reflect dietary pattern

#### 3.2.1 Richness and Shannon diversity are lower in plant-based than in omnivorous participants

Observed species richness and Shannon diversity differed significantly across the three dietary pattern groups (n=175; Kruskal-Wallis, p=0.0011 and p=0.0014, respectively), driven primarily by lower richness in vegans relative to omnivores (Dunn’s test, p.adj=0.001) and, to a lesser extent, in lacto-ovo-vegetarians relative to omnivores (p.adj=0.019); lacto-ovo-vegetarians and vegans did not differ from each other (p.adj=0.32) (Figure 2A). Shannon diversity followed the same pairwise pattern (Figure 2B). Pielou’s evenness did not differ significantly among groups (p=0.067) (Figure 2C).

**Figure 2.**
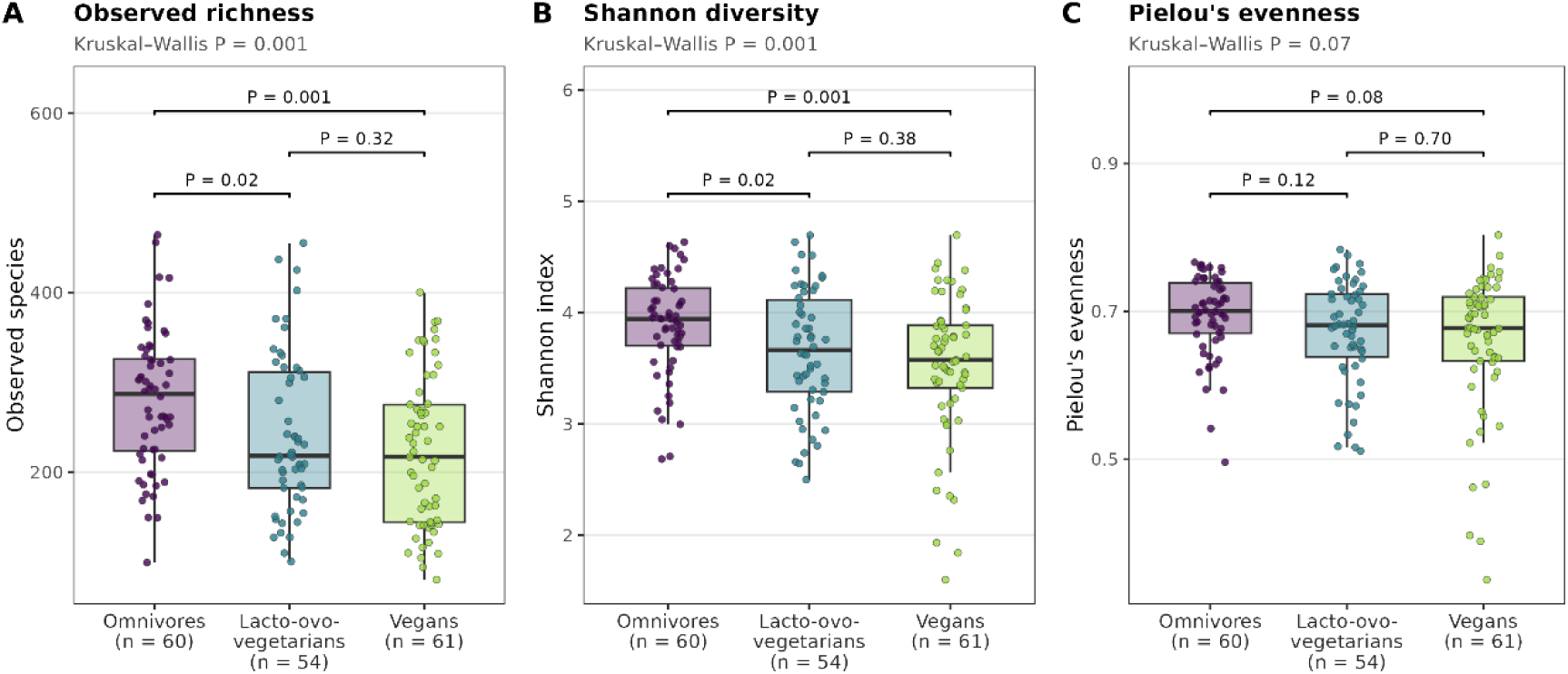
Alpha diversity of the gut microbiota by dietary pattern. (A) Observed species richness, (B) Shannon diversity and (C) Pielou’s evenness in omnivores (n = 60), lacto-ovo-vegetarians (n = 54) and vegans (n = 61). Boxes show the median and interquartile range, whiskers extend to 1.5 times the interquartile range, and points represent individual participants. Overall differences were assessed with the Kruskal–Wallis test (shown below each panel title), and pairwise comparisons with Dunn’s test with Benjamini–Hochberg correction (adjusted P values shown above the brackets). Alpha diversity was calculated on 2,249 species.

#### 3.2.2 Associations of plant-based diet indices with richness change after adjustment for dietary pattern

In partial Spearman correlations adjusted for sex, age, BMI and sequencing depth (n = 165), PDI and uPDI were negatively associated with richness (PDI: ρ = −0.216, 95% CI −0.359 to −0.063, q = 0.016; uPDI: ρ = −0.298, 95% CI −0.433 to −0.150, q = 0.001), whereas hPDI was not (ρ = −0.047, q = 0.66) (Figure 3). After additional adjustment for dietary pattern, the PDI association was no longer present (ρ = 0.016, q = 0.83), the uPDI association was attenuated but remained significant (ρ = −0.192, q = 0.020), and hPDI showed a positive association (ρ = 0.206, 95% CI 0.052 to 0.350, q = 0.016). Within individual dietary patterns, no index was associated with richness after correction for multiple comparisons. The direction of the hPDI association was positive in all three patterns (ρ = 0.17 to 0.24), and the largest uPDI estimate was observed in omnivores (ρ = −0.347, p = 0.013, q = 0.11).

**Figure 3.**
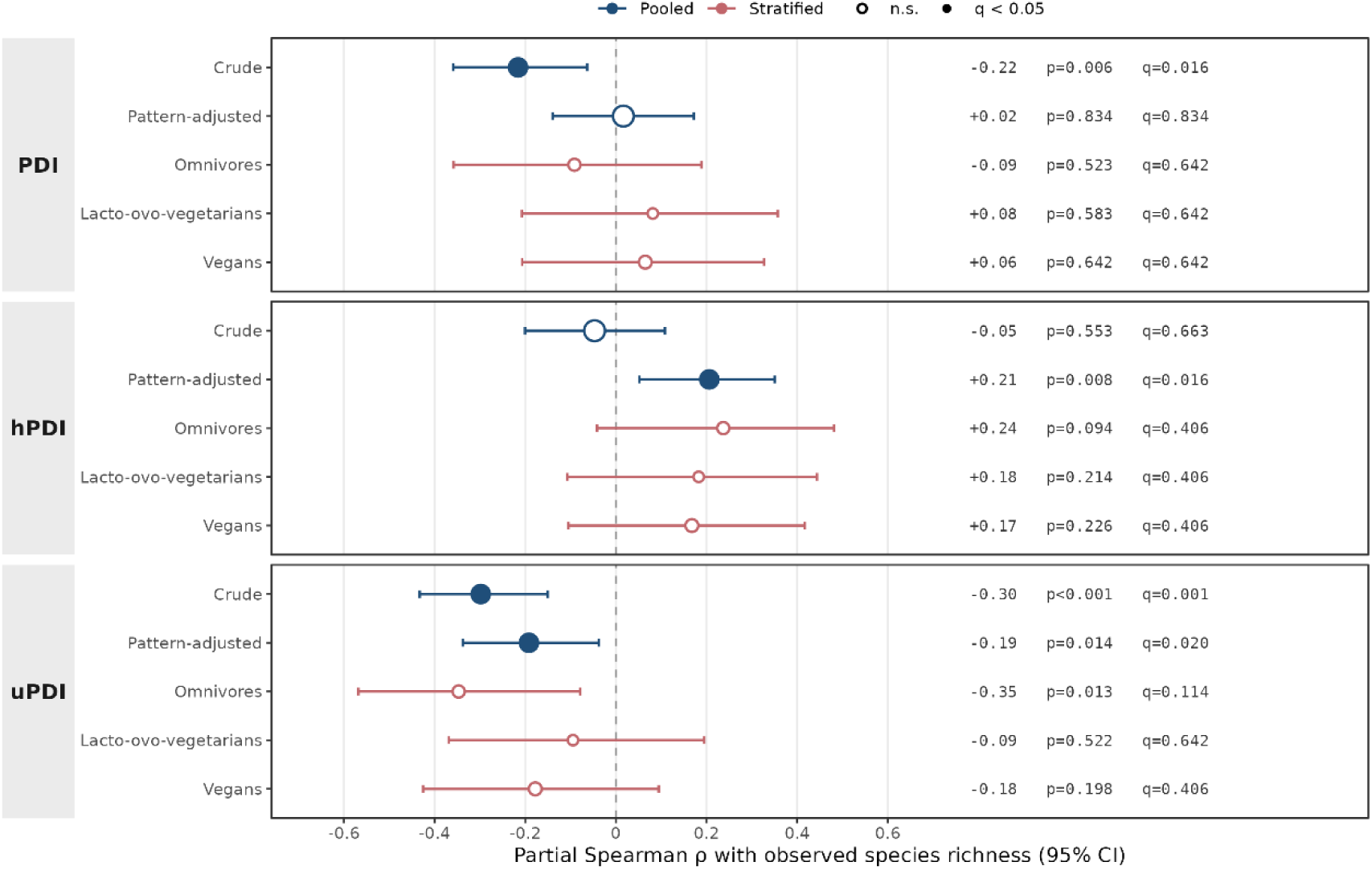
Associations between plant-based diet indices and observed species richness. Points show partial Spearman correlation coefficients with 95% confidence intervals (Fisher z transformation) for PDI, hPDI and uPDI, adjusted for sex, age, BMI and sequencing depth (n = 165; 55 omnivores, 52 lacto-ovo-vegetarians, 58 vegans). Blue: pooled analyses, unadjusted (crude) and additionally adjusted for dietary pattern (Spearman correlation of residuals after regressing both index and richness on dietary pattern and covariates). Red: analyses stratified by dietary pattern. Filled symbols indicate q < 0.05. Benjamini–Hochberg correction was applied separately to the pooled (6 tests) and stratified (9 tests) analyses.

#### 3.2.3 Plant-based diet indices vary mainly between dietary patterns, whereas MPPF intake varies mainly within them

Dietary pattern explained 62.0% of the variance in PDI and 42.9% in hPDI, compared with 15.7% for uPDI, 19.3% for hPDI_raw and 5.8% for MPPF intake (Figure 4). Within vegans, the coefficient of variation was 7.4% for PDI, 12.4% for hPDI and 11.6% for uPDI, compared with 22.5% for hPDI_raw and 26.8% for MPPF intake.

**Figure 4.**
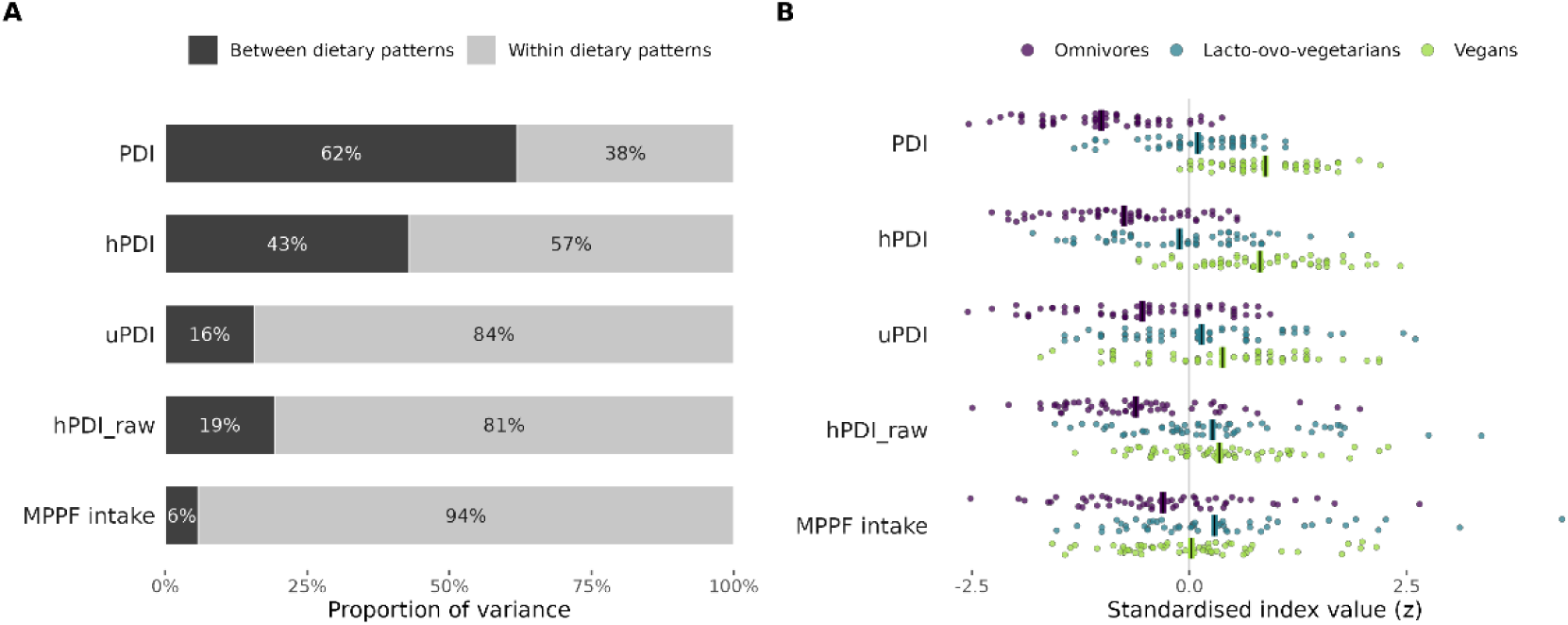
Between-and within-pattern variation of dietary indices (n = 165; 55 omnivores, 52 lacto-ovo-vegetarians, 58 vegans). (A) Proportion of the variance of each index explained by dietary pattern (between dietary patterns; η² from one-way analysis of variance) and remaining within dietary patterns. (B) Individual values of each index, standardised across the 165 participants, by dietary pattern; vertical bars indicate group means. hPDI_raw was computed from the same food groups as hPDI using continuous energy-adjusted intake instead of cohort-based quintiles.

### 3.3 MPPF intake is associated with higher gut microbial richness in plant-based but not omnivorous participants

In the analytic cohort (n = 172), MPPF intake ranged from 127 to 1,494 g/1,000 kcal (mean 636; 1 SD = 209 g/1,000 kcal). Mean values were 580 ± 207 g/1,000 kcal in omnivores, 689 ± 222 in lacto-ovo-vegetarians and 645 ± 188 in vegans (mean ± SD; Kruskal-Wallis p = 0.038), with most of its variance lying within rather than between dietary patterns (Figure 4; Table 1).

Plant-based participants derived a higher proportion of energy from ultra-processed foods than omnivores (adjusted difference 4.4 percentage points, 95% CI 1.1 to 7.6, p = 0.009; adjusted for sex, age and BMI), although differences across the three dietary patterns did not reach statistical significance (median 19.7%, 20.8% and 22.1% in omnivores, lacto-ovo-vegetarians and vegans, respectively; Kruskal-Wallis p = 0.061). Within each dietary pattern, MPPF intake was inversely correlated with the proportion of energy from ultra-processed foods (Spearman ρ = −0.42 in omnivores, −0.37 in lacto-ovo-vegetarians and −0.51 in vegans; all p ≤ 0.007).

In stratified models adjusted for sex, age, BMI and sequencing depth, MPPF intake was positively associated with richness among plant-based participants (β = 26.6 per SD, equivalent to 12.7 species per 100 g/1,000 kcal; 95% CI 10.9 to 42.3, p = 0.001, R² = 0.143) and was not associated among omnivores (β = −0.8, 95% CI −23.4 to 21.7, p = 0.94) (Figure 5A). The dietary pattern × MPPF intake interaction term was significant (β = 26.6 per SD, 95% CI 0.6 to 52.6, p = 0.045).

**Figure 5.**
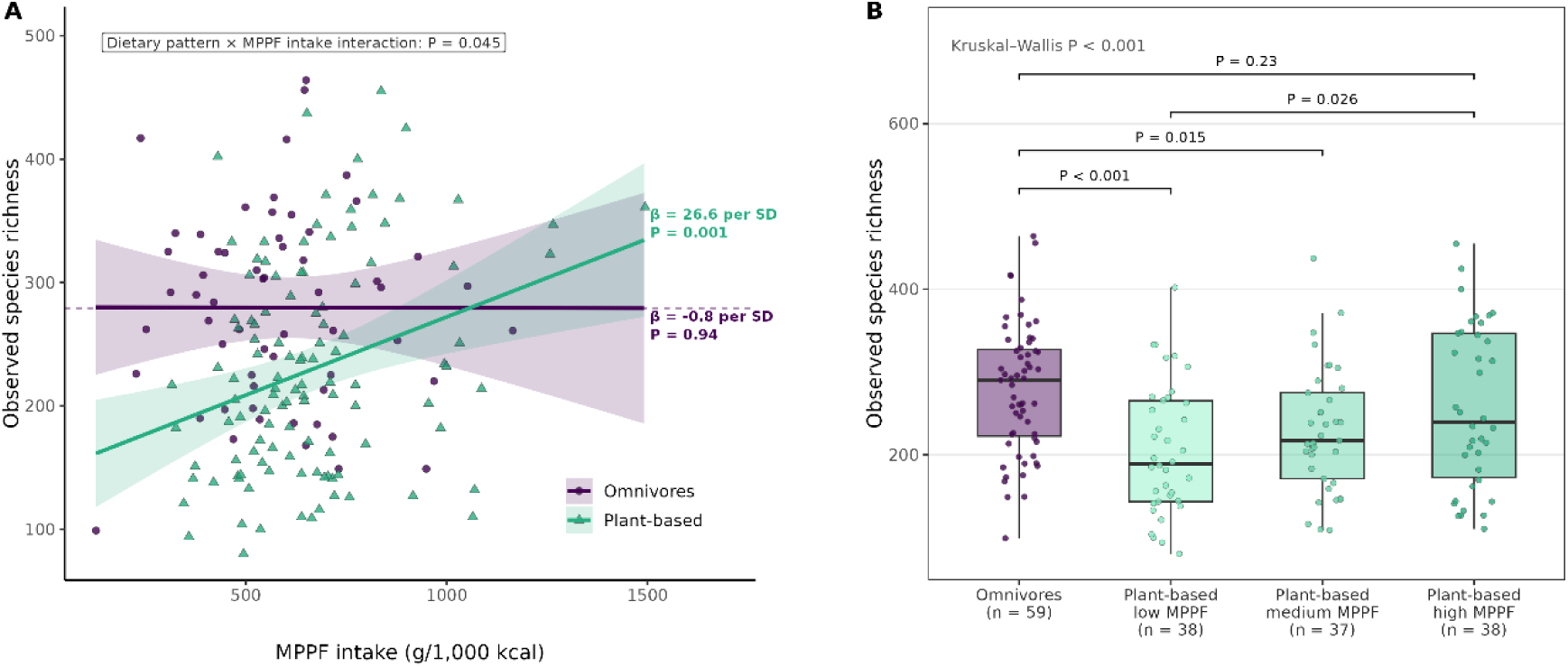
MPPF intake and observed species richness by dietary pattern. (A) Richness against MPPF intake in omnivores (n = 59) and plant-based participants (lacto-ovo-vegetarians and vegans combined; n = 113). Lines and shaded bands show predicted values and 95% confidence intervals from a linear model including a dietary pattern × MPPF intake interaction, adjusted for sex, age, BMI and sequencing depth; points are individual participants (circles, omnivores; triangles, plant-based participants). β values are within-group slopes per standard deviation of MPPF intake (209 g/1,000 kcal) from stratified models with the same covariates. The dashed line marks mean richness in omnivores. (B) Richness in omnivores and in plant-based participants divided into tertiles of MPPF intake (cut-offs 558 and 703 g/1,000 kcal). The overall difference was assessed with the Kruskal-Wallis test and pairwise comparisons with Dunn’s test with Benjamini-Hochberg correction; adjusted P values are shown for selected comparisons.

Within each of the three dietary patterns, the association was β = 27.7 per SD (95% CI 5.5 to 50.0, p = 0.016, R² = 0.265) in vegans, β = 22.7 (95% CI −0.6 to 46.0, p = 0.056) in lacto-ovo-vegetarians and β = −0.8 (p = 0.94) in omnivores; in this three-group model, the interaction terms relative to omnivores did not reach statistical significance (vegans, p = 0.055; lacto-ovo-vegetarians, p = 0.14).

Plant-based participants were divided into tertiles of MPPF intake (cut-offs 558 and 703 g/1,000 kcal) and compared with omnivores (Kruskal-Wallis χ² = 19.1, p < 0.001) (Figure 5B). Median richness was 290 species in omnivores and 189, 217 and 239 species in the low, medium and high MPPF intake tertiles, respectively. Richness was lower than in omnivores in the low (p.adj < 0.001) and medium (p.adj = 0.015) tertiles but not in the high tertile (p.adj = 0.23), and was higher in the high than in the low tertile (p.adj = 0.026).

The association between MPPF intake and richness within plant-based participants was robust across sensitivity analyses (Supplementary Table S3; Supplementary Fig. S1). It was unchanged when MPPF intake included the five items whose processing level depends on commercial formulation (54-item version; β = 27.3 per SD, p = 0.001), when intake was expressed as servings instead of grams per 1,000 kcal (β = 26.9, p = 0.003) and when estimated by robust regression (Huber M-estimation; β = 28.2, p < 0.001). It remained significant when each of the 49 MPPF items (β range 23.7 to 28.3; all p ≤ 0.003) and each participant (β range 24.5 to 30.0; all p ≤ 0.004) was omitted in turn. When richness was defined using increasing relative-abundance detection thresholds, the association remained significant at 0.001% (β = 24.6, p = 0.001), 0.01% (β = 18.7, p < 0.001) and 0.1% (β = 6.1, p = 0.008), corresponding to 11.4%, 11.5% and 7.7% of mean richness per SD, but not at 1% (β = 0.3, p = 0.52), at which mean richness was 19 species. Among omnivores, no association was observed in any of these analyses (β range −7.5 to 7.5 per SD; all p ≥ 0.45). For the dietary pattern × MPPF intake interaction, robust regression yielded a larger estimate than ordinary least squares (β = 32.8 vs 26.6 per SD; p = 0.015 vs 0.045), and estimates in the 54-item, servings-based and leave-one-out analyses ranged from 21.5 to 34.2 per SD; given the baseline p value of 0.045, p exceeded 0.05 when some items (13 of 49) or participants (41 of 172) were omitted.

The association was also independent of fibre intake. In the 158 participants with 3-day dietary records, an instrument independent of the questionnaire used to derive MPPF intake, the association among plant-based participants was 27.3 per SD (p = 0.002) before and 26.9 per SD (95% CI 9.6 to 44.1, p = 0.003) after additional adjustment for fibre intake estimated from the records, and the dietary pattern × MPPF intake interaction estimate was likewise unchanged (26.6 and 26.7 per SD) (Supplementary Table S3; Supplementary Fig. S1).

### 3.4 Gut microbial composition differs by dietary pattern and varies with MPPF intake only in plant-based participants

Dietary pattern was associated with overall microbial composition (Aitchison distance, PERMANOVA, R² = 0.023, p < 0.001), with consistent results using rclr (R² = 0.044), Jaccard (R² = 0.031) and Bray-Curtis (R² = 0.025) distances (all p ≤ 0.001) (Figure 6A). All pairwise comparisons between dietary patterns were significant (Aitchison, BH-adjusted p ≤ 0.011). Multivariate dispersion differed among dietary patterns (PERMDISP, F = 3.86, p = 0.020), with lower dispersion in vegans than in omnivores (Tukey, p = 0.028).

**Figure 6.**
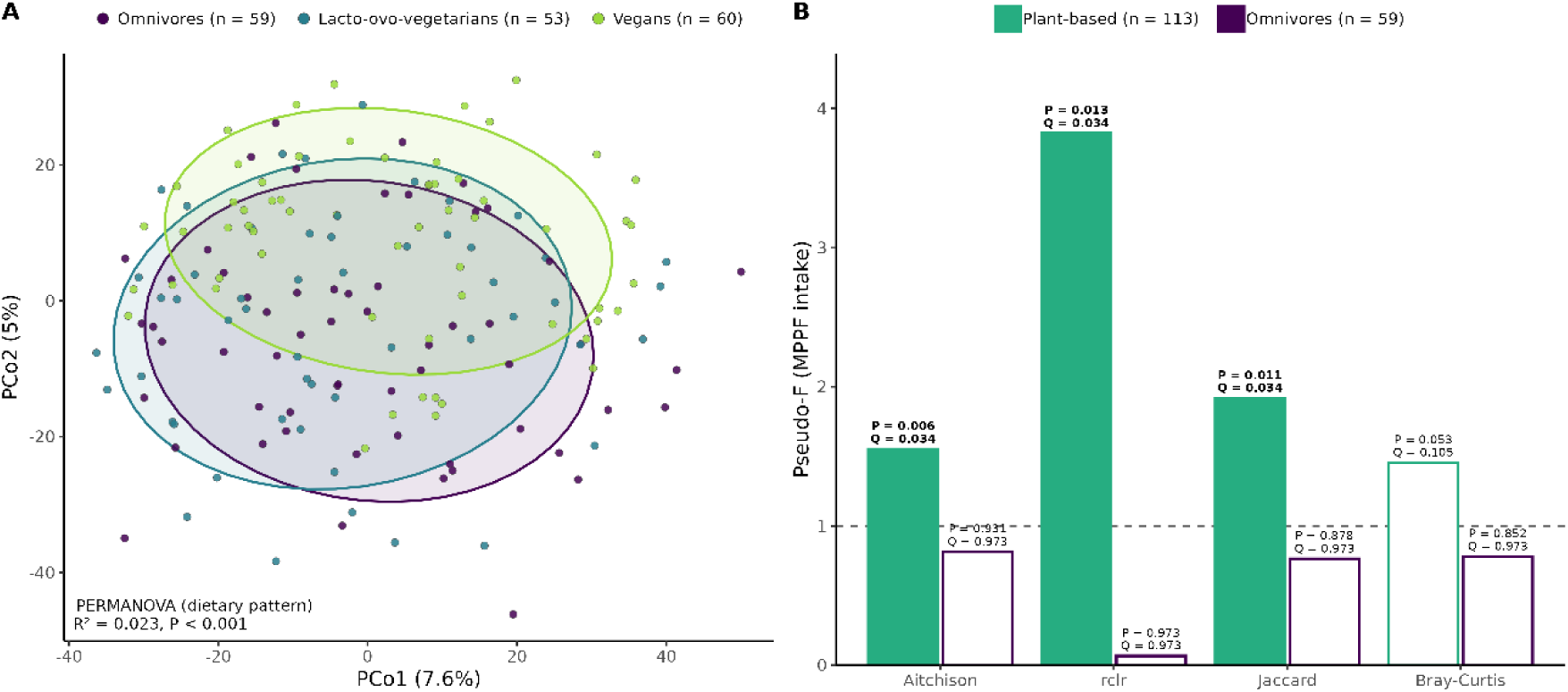
Gut microbial community composition by dietary pattern and MPPF intake. (A) Principal coordinates analysis (PCoA) based on Aitchison distance. Each point represents one participant; ellipses represent one standard deviation (68% normal-theory) around each dietary pattern centroid. PERMANOVA was adjusted for MPPF intake, sex, age and BMI. (B) PERMANOVA pseudo-F for MPPF intake within plant-based participants and omnivores across four distance metrics, adjusted for sex, age and BMI. Pseudo-F values above the dashed line (F = 1) indicate that MPPF intake explained more compositional variation than expected by chance. Q values were obtained with Benjamini-Hochberg correction across the eight stratified tests (2 strata × 4 metrics). Filled bars indicate q < 0.05; open bars indicate non-significant results. All PERMANOVA analyses used 9,999 permutations. rclr, robust centred log-ratio.

In the full cohort, adjusting for dietary pattern, sex, age and BMI, MPPF intake was associated with microbial composition using Aitchison (R² = 0.008, p = 0.022) and Jaccard (R² = 0.009, p = 0.027) distances, but not using rclr (R² = 0.003, p = 0.64) or Bray-Curtis (R² = 0.008, p = 0.061). Within plant-based participants (n = 113), MPPF intake was associated with microbial composition using Aitchison (R² = 0.014, p = 0.006), rclr (R² = 0.033, p = 0.013) and Jaccard (R² = 0.017, p = 0.011) distances (all q = 0.034), but not using Bray-Curtis (R² = 0.013, p = 0.053, q = 0.105). Within omnivores (n = 59), no association was observed with any metric (all p ≥ 0.85, q = 0.97) (Figure 6B; Supplementary Table S4).

### 3.5 MPPF intake is associated with more health-favourable gut species in plant-based participants

All 50 species ranked most favourably and 41 of the 50 ranked most unfavourably for cardiometabolic health in the ZOE Microbiome Health Ranking 2025 ^60^ were detected among plant-based participants (mean 19.7 and 9.5 species per participant, respectively). Within plant-based participants, MPPF intake was positively associated with the number of health-favourable species (14.6% per SD, 95% CI 3.5 to 26.8, p = 0.010) and was not associated with the number of health-unfavourable species (−7.0% per SD, 95% CI −15.6 to 2.6, p = 0.15) (Table 2). Prevalence-matched random species sets increased by 6% and 8% per SD, respectively; relative to these sets, the number of health-unfavourable species increased less than expected (p = 0.022), and the number of health-favourable species increased more than expected without reaching statistical significance (p = 0.060). MPPF intake was also associated with the number of diet-favourable species (11.5% per SD, 95% CI 3.3 to 20.4, p = 0.006), although this association did not exceed that of random sets (p = 0.17). Within omnivores, no association was observed for any set (all p ≥ 0.40).

**Table 2.**
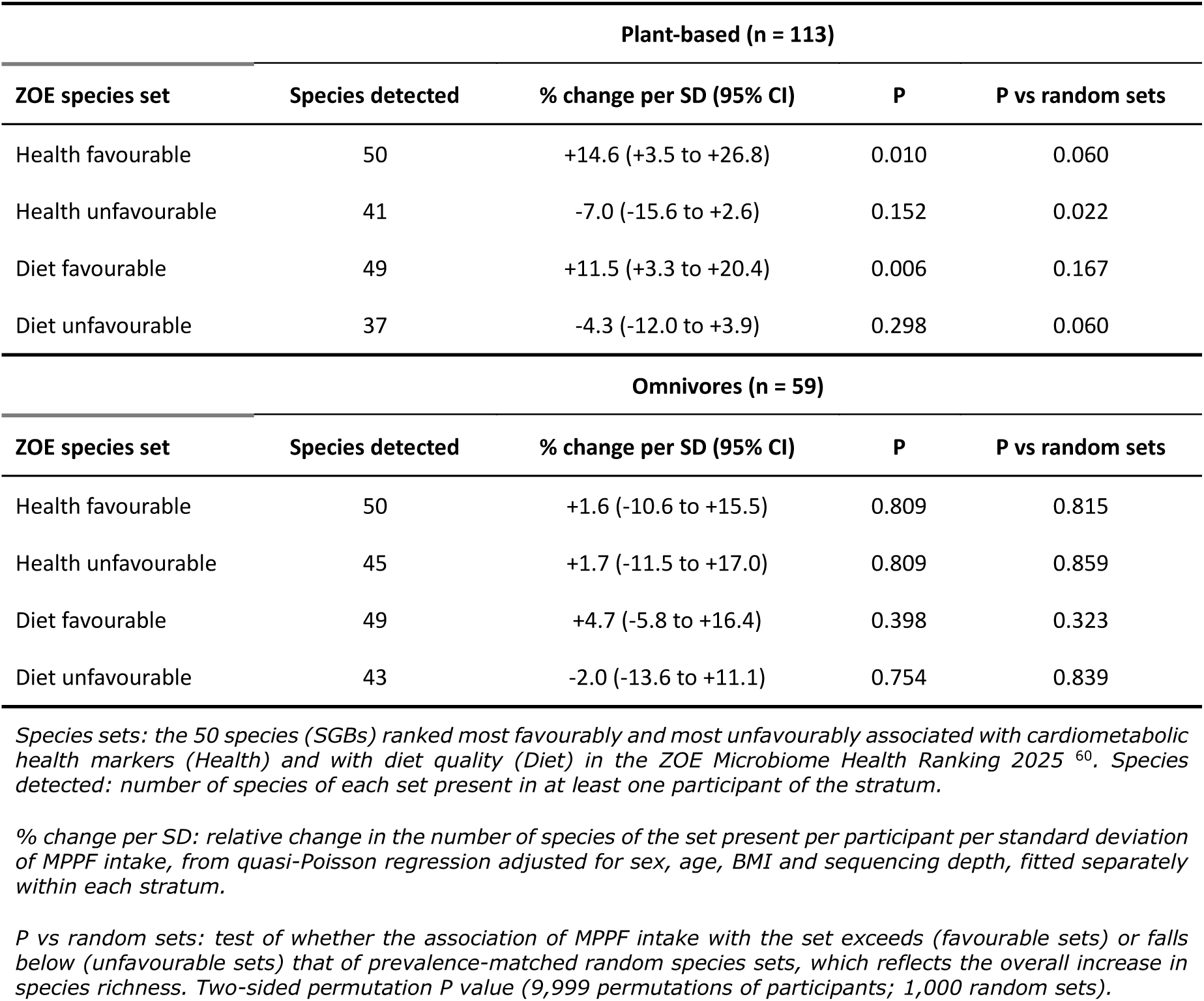
Association between MPPF intake and the number of ZOE health-and diet-ranked gut species, by dietary stratum.

|  | **Plant-based (n = 113)** |  |  |  |
|---|---|---|---|---|
| **ZOE species set** | **Species detected** | **% change per SD (95% CI)** | **P** | **P vs random sets** |
| Health favourable | 50 | +14.6 (+3.5 to +26.8) | 0.010 | 0.060 |
| Health unfavourable | 41 | -7.0 (-15.6 to +2.6) | 0.152 | 0.022 |
| Diet favourable | 49 | +11.5 (+3.3 to +20.4) | 0.006 | 0.167 |
| Diet unfavourable | 37 | -4.3 (-12.0 to +3.9) | 0.298 | 0.060 |
|  | **Omnivores (n = 59)** |  |  |  |
| **ZOE species set** | **Species detected** | **% change per SD (95% CI)** | **P** | **P vs random sets** |
| Health favourable | 50 | +1.6 (-10.6 to +15.5) | 0.809 | 0.815 |
| Health unfavourable | 45 | +1.7 (-11.5 to +17.0) | 0.809 | 0.859 |
| Diet favourable | 49 | +4.7 (-5.8 to +16.4) | 0.398 | 0.323 |
| Diet unfavourable | 43 | -2.0 (-13.6 to +11.1) | 0.754 | 0.839 |

*Species sets: the 50 species (SGBs) ranked most favourably and most unfavourably associated with cardiometabolic health markers (Health) and with diet quality (Diet) in the ZOE Microbiome Health Ranking 2025 <sup>60</sup>. Species detected: number of species of each set present in at least one participant of the stratum.*
*% change per SD: relative change in the number of species of the set present per participant per standard deviation of MPPF intake, from quasi-Poisson regression adjusted for sex, age, BMI and sequencing depth, fitted separately within each stratum.*
*P vs random sets: test of whether the association of MPPF intake with the set exceeds (favourable sets) or falls below (unfavourable sets) that of prevalence-matched random species sets, which reflects the overall increase in species richness. Two-sided permutation P value (9,999 permutations of participants; 1,000 random sets).*

The association with health-favourable species was similar when the 100 most favourably ranked species were used (13.9% per SD, p = 0.006), when species presence was defined at relative-abundance thresholds of 0.001% and 0.01%, after additional adjustment for vegan versus lacto-ovo-vegetarian pattern, and when each participant was omitted in turn (all p ≤ 0.021).

At the species level (exploratory; Supplementary Table S5), 40 of the 46 testable health-favourable species were more frequently detected at higher MPPF intake among plant-based participants, 15 of them with nominal p < 0.05, including Vescimonas sp000435555 (Oscillospiraceae; OR 2.13 per SD, 95% CI 1.33 to 3.43, p = 0.002, q = 0.080) and three CAG-170 species (OR 1.57 to 1.69, p = 0.018 to 0.041). Among health-unfavourable species, Ruminococcus_B gnavus (OR 0.54, 95% CI 0.32 to 0.91, p = 0.021) and Blautia_A fusiformis (OR 0.53, 95% CI 0.34 to 0.84, p = 0.007) were less frequently detected at higher MPPF intake. No species-level association remained significant after correction for multiple testing.

### 3.6 Self-reported health is better at higher MPPF intake in plant-based participants, with no differences in biochemical parameters

The association between MPPF intake and SF-36 total score differed between omnivores and the combined plant-based group (dietary pattern × MPPF intake interaction, β = 3.8 per SD, 95% CI 0.6 to 7.0, p = 0.022). Within the plant-based group (n = 113), MPPF intake was positively associated with SF-36 total score (β = 3.0 per SD, 95% CI 1.1 to 4.9, p = 0.002, R² = 0.123), whereas no association was observed within omnivores (n = 59; β = −1.2, 95% CI −4.1 to 1.8, p = 0.43) (Supplementary Fig. S2).

When the three dietary patterns were modelled separately, within-group estimates were similar in lacto-ovo-vegetarians (n = 53; β = 3.4 per SD, p = 0.020, R² = 0.143) and vegans (n = 60; β = 2.8, p = 0.037, R² = 0.131).

Species richness was not correlated with SF-36 total score (Spearman ρ = 0.096, p = 0.21).

MPPF intake was not associated with any biochemical or anthropometric parameter, either in the full cohort or within dietary patterns (all q ≥ 0.22). PDI, hPDI and uPDI were associated with several biochemical parameters without adjustment for dietary pattern, mainly lipid and leukocyte measures that also differed between patterns (Table 1), but none of these associations remained after adjustment for dietary pattern (all q ≥ 0.51).

## 4. Discussion

In this cross-sectional study, dietary pattern was associated with gut microbial richness and with overall community composition, with lower richness in participants who avoided animal foods. Established plant-based diet indices largely reflected dietary pattern itself and retained little variation within patterns. Within plant-based participants, the energy-adjusted intake of minimally processed plant foods was associated with higher species richness, with community composition, with a higher number of species previously ranked as favourably associated with cardiometabolic health, and with better self-reported health. None of these associations was observed among omnivores.

The lower richness we observed in vegans and lacto-ovo-vegetarians relative to omnivores is consistent with the largest metagenomic comparison published to date ^14^ and contrasts with smaller studies reporting higher richness in vegetarians ^12^ or no difference between vegans and omnivores ^13^. Part of this inconsistency is methodological: studies differ in whether they use 16S rRNA gene sequencing or shotgun sequencing, in profiling tools and reference databases, in sample size, and in how vegetarian and vegan diets are defined, usually by self-report ^9,11^. The inconsistency may also be conceptual: a dietary label states which foods are absent, not which foods take their place, and people following the same pattern may eat very differently ^4^. Comparisons between labels may therefore involve groups whose diets vary as much within as between them, a possibility that our data allowed us to examine directly.

Plant-based diet indices were not designed to answer the question we asked of them. PDI, hPDI and uPDI were developed to predict type 2 diabetes and coronary heart disease in predominantly omnivorous US cohorts, and they perform well for that purpose ^15,16^; the limitation we describe is one of scope rather than of quality. Food-group intakes are scored in quintiles of each cohort’s own distribution, an approach with recognised constraints ^23^, and participants who already avoid animal foods are compressed into the upper part of that range. Our data quantify this: dietary pattern accounted for 62.0% of the variance in PDI and 42.9% in hPDI, against 5.8% for MPPF intake, and within vegans the coefficient of variation was 7.4% for PDI and 12.4% for hPDI, against 26.8% for MPPF intake. The consequence is visible in the hPDI results: no association with richness appeared in the pooled analysis, and a positive one emerged only after adjustment for dietary pattern, because vegans score highest on hPDI yet had the lowest richness, so that pattern-level and within-pattern effects ran in opposite directions.

This leads to the main implication of our findings: comparing microbial diversity between dietary labels without characterising what is eaten within them may be misleading. Among plant-based participants, MPPF intake was associated with richness (26.6 species per standard deviation of MPPF intake, equivalent to 12.7 species per 100 g/1,000 kcal; p = 0.001), and the association held under a broader MPPF definition, robust regression, the omission of each item and each participant in turn, and additional adjustment for fibre intake measured with 3-day dietary records. Among omnivores there was no association in any of these analyses. When plant-based participants were grouped by tertiles of MPPF intake, richness was lower than in omnivores in the low and medium tertiles but not in the high tertile, where the median was 239 species against 290 in omnivores. If this pattern holds, the average richness reported for a plant-based group may depend largely on how much of its members’ diets consists of minimally processed plant foods. Between-label comparisons would then reflect the dietary make-up of each particular sample, rather than an effect of excluding animal foods per se. This is not a remote possibility: plant-based participants in our cohort derived a higher proportion of energy from ultra-processed foods than omnivores, as reported in NutriNet-Santé ^4^ and UK Biobank ^5^. One explanation is that when animal foods are excluded, plant foods become almost the only source of fermentable substrate reaching the colon ^7,8^, so that variation in their quantity and structural integrity carries more weight. Although the formal dietary pattern by MPPF intake interaction was of borderline significance (p = 0.045), the within-group association was consistent across all sensitivity analyses among plant-based participants and absent among omnivores.

The signal appears to lie among species present at low abundance. The association with richness persisted at detection thresholds up to 0.1% relative abundance but disappeared at 1%, by which point fewer than twenty species per sample remained. The composition results point the same way: within plant-based participants MPPF intake was associated with Aitchison, rclr and Jaccard distances, which give comparatively more weight to rare taxa and to presence or absence, but not with Bray-Curtis, which is dominated by abundant taxa. Higher intakes of minimally processed plant foods were therefore associated with the detection of additional low-abundance species rather than with a rearrangement of the dominant members of the community.

Which species are detected matters as much as how many. Using an externally defined ranking derived from more than 34,000 individuals ^60^, the number of species ranked as favourably associated with cardiometabolic health was higher at higher MPPF intake among plant-based participants, whereas the number of unfavourably ranked species was not. Because richness itself increased, we compared these sets with prevalence-matched random sets: unfavourable species increased less than expected from the overall gain in richness (p = 0.022), and favourable species showed the opposite tendency (p = 0.060). In exploratory species-level analyses, the species more frequently detected at higher MPPF intake were largely uncultured Clostridia, including Vescimonas sp000435555 and three species of CAG-170. CAG-170 was recently identified as the strongest health-associated gut lineage across 11,115 metagenomes from 39 countries, with predicted capacity for vitamin B12 biosynthesis and cross-feeding ^61^, and its diversity has been inversely associated with body mass index ^62^. Ruminococcus_B gnavus and Blautia_A fusiformis, both unfavourably ranked, were less frequently detected at higher MPPF intake; the former is associated with Crohn’s disease and produces a glucorhamnan polysaccharide that induces TNF-α secretion ^63^. No species-level result survived correction for multiple testing.

Calling a food healthful requires a criterion, and the criterion determines what the resulting score can detect. In the plant-based diet indices, food groups were assembled by nutrient and culinary similarity and classified as healthful or unhealthful according to prior associations with type 2 diabetes, cardiovascular disease and certain cancers ^15^. Foods that protect against cardiometabolic disease are not necessarily those that matter most for the gut microbiota. Processing operates along a different axis, altering the food matrix and its microstructure, and thereby which components reach the colon and in what form ^25,64^. A single hPDI category can span very different processing levels: whole grains include intact grains and ultra-processed breakfast cereals; nuts include raw and roasted salted products; legumes include cooked pulses alongside tofu, tempeh and textured soy protein; and vegetable oils are scored as healthful although they contain neither fibre nor food matrix. The converse also holds, since boiled potatoes, white rice and freshly squeezed juice are minimally processed yet scored as less healthful. Plant-based meat and dairy analogues, typically ultra-processed, have no predefined place in the indices at all. Our data support this distinction: the association between MPPF intake and richness was unchanged after adjustment for fibre intake measured with 3-day dietary records, an instrument independent of the questionnaire, indicating that what MPPF intake captures is not reducible to the amount of fibre consumed. Within each dietary pattern, MPPF intake was inversely correlated with the proportion of energy from ultra-processed foods (ρ = −0.37 to −0.51), as expected given that both draw on the same energy budget; our design cannot separate eating more minimally processed foods from eating fewer ultra-processed products, whose additives may themselves affect the gut microbiota ^26^.

MPPF intake was also associated with self-reported health among plant-based participants and not among omnivores, mirroring the richness results. Richness itself was not correlated with the SF-36 total score, so the microbiome does not appear to account for this association. Two hypotheses may be considered. The first is that within plant-based diets MPPF intake separates different profiles of adherents, for example those motivated by health and whole-food eating from those relying more on processed plant-based products, whereas among omnivores health-oriented habits may be expressed through dietary components that MPPF intake does not capture ^65,66^. Health-motivated vegans, for instance, report eating more fruit and fewer sweets than those motivated by ethical concerns ^65^. The second is the substitution argument advanced above: when minimally processed plant foods form the core of the diet, replacing them with processed products may have consequences that animal foods buffer in omnivorous diets. Both hypotheses remain to be tested, and the SF-36 is a self-reported measure.

The study has several strengths. Diet was characterised with a food frequency questionnaire extended to include plant-based foods and products, and every item was classified individually according to NOVA, with the full item-level classification and its justification provided in the supplementary material. Energy and fibre intake were also assessed with 3-day dietary records, an instrument independent of the vFFQ, which allowed the main association to be tested with adjustment for fibre from a separate source. The microbiome was profiled by shotgun metagenomics at species level rather than by 16S rRNA gene sequencing. The three dietary patterns were recruited in comparable numbers, which gives reasonable resolution within as well as between patterns. The main association was examined under alternative MPPF definitions, estimators and detection thresholds, and with individual items and participants omitted. The analysis of health-associated species used an external ranking defined a priori in an independent and much larger cohort, with prevalence-matched random sets as a comparison.

Several limitations should be considered. The design is cross-sectional, so no causal or directional inference is possible. The extended questionnaire has not yet been validated, with a validation study underway, and dietary assessment by recall is subject to measurement error. Items whose processing level depends on commercial formulation were classified by their predominant presentation in the Spanish market, a decision mitigated but not resolved by the broader 54-item sensitivity analysis. Participants were young and healthy, with a narrow range of body mass index, and recruited in a single city, which limits generalisability. Statistical power was limited for interaction terms and for individual taxa, and several analyses were exploratory.

In summary, dietary labels alone appear to be an insufficient description of diet in gut microbiome research, and the degree of processing of the plant foods consumed is a dimension worth measuring explicitly rather than assuming. Among people who avoid animal foods, those whose diets were built on minimally processed plant foods had a richer gut microbiota, containing more of the species independently associated with cardiometabolic health, than those whose diets contained fewer minimally processed plant foods. Establishing whether this reflects an effect of the foods themselves will require validation of the extended questionnaire, longitudinal studies and dietary interventions that manipulate the processing level of plant foods while holding dietary pattern constant.

## Data availability

The raw shotgun metagenomic sequencing reads generated in this study have been deposited in the European Nucleotide Archive (ENA) and are currently held under private access. The data and corresponding accession number will be made publicly available upon publication of the peer-reviewed article. Individual-level dietary, clinical and questionnaire data are not publicly available because they contain information that could compromise participant privacy under the consent obtained; de-identified data are available from the corresponding author upon reasonable request and subject to approval by the Research Ethics Committee of Fundación IMDEA Nutrición.

## Code availability

All code used for data processing, statistical analysis and figure generation is maintained in a private GitHub repository and will be made publicly available upon publication of the peer-reviewed article. Until then, the code is available from the corresponding author upon reasonable request.

## Acknowledgements

We thank all study participants for their time and commitment, and ProVeg Spain and the Unión Vegetariana Española (UVE) for their contribution to data collection.

## Author contributions

A.M.-H. and L.J.M.-Z. conceived the study. A.A., H.M.-P., A.C., A.M., E.A.-A., A.B.-R., A.P.-C. and L.J.M.-Z. recruited participants and conducted the study visits. A.A., H.M.-P., A.C., A.M. and E.A.-A. carried out the dietary assessment, including the 3-day dietary records, the review of portion sizes and food composition, and the classification of food items into NOVA groups and plant-based diet index food groups. D.C.-C. contributed to the dietary and metagenomic methodology. A.B.-R. led the faecal sample processing and DNA extraction, with contributions from B.L.-P., A.P.-C. and N.C.-R. A.M.-H. curated and processed the dietary and metagenomic data, developed the analysis code, performed the statistical analyses, prepared the figures and wrote the first draft of the manuscript. L.J.M.-Z. acquired funding and administered the project, and L.J.M.-Z. and E.A.-A. supervised the work. All authors contributed to the interpretation of the results, critically reviewed the manuscript and approved the final version.

