## Supplementary Information (Tables S2-S5, Figures S1-S2) for "Not all plant-based diets are alike: intake of minimally processed plant foods is associated with gut microbial richness among vegetarians and vegans"

### Contents


| | | |
|---|---|---|
| **Supplementary Table S1** | Vegan-adapted food frequency questionnaire (vFFQ): items, portions and energy, plant-based diet index food groups and NOVA classification | Separate Excel file |
| **Supplementary Table S2** | Remaining biochemical and haematological parameters by dietary pattern | 2 |
| **Supplementary Table S3** | Sensitivity analyses of the association between MPPF intake and observed species richness | 3 |
| **Supplementary Table S4** | Association between MPPF intake and microbiome composition, overall and within dietary strata | 4 |
| **Supplementary Table S5** | Species of the ZOE health ranking and their association with MPPF intake, by dietary stratum | 6 |
| **Supplementary Fig. S1** | Sensitivity analyses of the association between MPPF intake and observed species richness | 12 |
| **Supplementary Fig. S2** | MPPF intake and SF-36 total score by dietary pattern | 13 |

Supplementary Table S2. Remaining biochemical and haematological parameters by dietary pattern.

| Characteristic | Omnivores (n=59) | Lacto-ovo-vegetarians (n=53) | Vegans (n=60) | p | q |
|---|---|---|---|---|---|
| Red blood cells, $\times 10^6/\mu L$ | 4.8 (0.4) | 4.7 (0.4) | 4.6 (0.4) | 0.012 | 0.032 |
| MCH, pg | 29.9 (1.5) | 30.1 (1.8) | 29.9 (2.5) | 0.393 | 0.596 |
| MCHC, g/dL | 33.0 (0.5) | 33.1 (0.6) | 33.1 (0.8) | 0.426 | 0.623 |
| Platelets, $\times 10^3/\mu L$ | 236 (58) | 240 (46) | 222 (49) | 0.176 | 0.301 |
| MPV, fL | 9.3 (1.0) | 9.5 (1.0) | 10.0 (1.1) | <0.001 | 0.004 |
| Lymphocytes, % | 36.1 (7.4) | 33.1 (7.7) | 34.0 (9.1) | 0.093 | 0.201 |
| Monocytes, % | 8.0 (1.5) | 8.3 (2.0) | 8.4 (2.0) | 0.681 | 0.822 |
| Neutrophils, % | 52.5 (8.2) | 55.4 (8.6) | 54.5 (8.8) | 0.151 | 0.291 |
| Eosinophils, % | 2.6 (1.9) | 2.5 (1.7) | 2.3 (1.4) | 0.695 | 0.822 |
| Basophils, % | 0.7 (0.4) | 0.8 (0.4) | 0.8 (0.4) | 0.592 | 0.809 |
| Monocytes, cells/µL | 487 (141) | 446 (115) | 415 (170) | 0.004 | 0.013 |
| Basophils, cells/µL | 45 (24) | 42 (18) | 38 (18) | 0.287 | 0.470 |
| Eosinophils, cells/µL | 163 (135) | 137 (92) | 114 (77) | 0.088 | 0.201 |
| Neutrophils, cells/µL | 3284 (1321) | 3119 (1237) | 2757 (1175) | 0.009 | 0.027 |
| Magnesium, mmol/L | 0.86 (0.05) | 0.85 (0.05) | 0.85 (0.04) | 0.442 | 0.625 |
| Phosphorus, mg/dL | 3.4 (0.4) | 3.3 (0.5) | 3.2 (0.4) | 0.103 | 0.212 |
| Total cholesterol / HDL | 2.96 (0.66) | 2.65 (0.63) | 2.64 (0.56) | 0.005 | 0.015 |
| LDL / HDL | 1.74 (0.56) | 1.41 (0.54) | 1.39 (0.47) | <0.001 | 0.003 |
| Triglycerides / HDL | 1.12 (0.68) | 1.19 (0.82) | 1.21 (0.64) | 0.348 | 0.548 |

*Values are mean (SD). p-values from Kruskal-Wallis tests; q-values are Benjamini-Hochberg adjusted across all 41 biochemical parameters measured, the same family used in Table 1. MCH, mean corpuscular haemoglobin; MCHC, mean corpuscular haemoglobin concentration; MPV, mean platelet volume.*

Supplementary Table S3. Sensitivity analyses of the association between MPPF intake and observed species richness.

| Analysis | Plant-based (n = 113) | Omnivores (n = 59) | Pattern × MPPF intake interaction (n = 172) |
|---|---|---|---|
| Primary analysis (49 items) | 26.6 (0.001) | −0.8 (0.941) | 26.6 (0.045) |
| 54-item version | 27.3 (0.001) | 0.1 (0.993) | 26.4 (0.0495) |
| MPPF intake in servings/1,000 kcal | 26.9 (0.003) | 1.4 (0.895) | 25.1 (0.052) |
| Robust regression (Huber M-estimation) | 28.2 (<0.001) | −2.6 (0.815) | 32.8 (0.015) |
| Robust regression (MM-estimation) | 28.2 (<0.001) | −3.1 (0.790) | 32.6 (0.017) |
| Leave-one-out, MPPF items | 23.7 to 28.3; 49/49 with p < 0.05 | −4.6 to 7.5; 0/49 with p < 0.05 | 21.5 to 28.5; 36/49 with p < 0.05 |
| Leave-one-out, participants | 24.5 to 30.0; 113/113 with p < 0.05 | −7.5 to 4.0; 0/59 with p < 0.05 | 22.4 to 34.2; 131/172 with p < 0.05 |
| Detection threshold 0.001% | 24.6 (0.001) | −1.1 (0.914) | 24.9 (0.042) |
| Detection threshold 0.01% | 18.7 (<0.001) | −0.5 (0.953) | 19.1 (0.031) |
| Detection threshold 0.1% | 6.1 (0.008) | 0.5 (0.864) | 5.3 (0.158) |
| Detection threshold 1% | 0.3 (0.521) | 0.4 (0.448) | 0.0 (0.977) |
| Participants with 3-day dietary records | 27.3 (0.002) | −1.4 (0.902) | 26.6 (0.054) |
| + fibre intake from 3-day dietary records | 26.9 (0.003) | −3.0 (0.794) | 26.7 (0.054) |

Values are β per SD of MPPF intake (209 g/1,000 kcal) with p values in parentheses, from linear models of observed species richness adjusted for sex, age, BMI and sequencing depth. For leave-one-out analyses, the range of β across iterations and the number of iterations with $p < 0.05$ are shown (49 items; 113, 59 and 172 participants, respectively).

MPPF intake in servings/1,000 kcal: the same 49 items, with FFQ frequency categories converted to servings per day instead of grams.

54-item version: primary measure plus five items whose processing level depends on commercial formulation (tofu, tempeh, natto, roasted peanuts and textured soy protein).

Robust regression: Huber M-estimation (MASS::rlm; p values approximated from t statistics) and MM-estimation (robustbase::lmrob; p values as reported) as a check.

Detection thresholds: richness recomputed counting only species above the stated relative abundance in each sample.

3-day dietary records: participants with records available (n = 158; 100 plant-based, 58 omnivores); the primary model was refitted in this subsample and then additionally adjusted for total fibre intake (g/1,000 kcal) estimated from the records.

Supplementary Table S4. Association between MPPF intake and microbiome composition, overall and within dietary strata.

| Analysis | n | $R^2$ (%) | Pseudo-F | P | Q |
|---|---|---|---|---|---|
| **Aitchison** | | | | | |
| Full cohort (pattern-adjusted) | 172 | 0.79 | 1.39 | 0.022 | – |
| Plant-based | 113 | 1.36 | 1.55 | 0.006 | 0.034 |
| *Vegans* | 60 | 2.14 | 1.28 | 0.049 | – |
| *Lacto-ovo-vegetarians* | 53 | 2.29 | 1.22 | 0.088 | – |
| Omnivores | 59 | 1.40 | 0.81 | 0.931 | 0.973 |
| **rclr** | | | | | |
| Full cohort (pattern-adjusted) | 172 | 0.31 | 0.55 | 0.643 | – |
| Plant-based | 113 | 3.25 | 3.82 | 0.013 | 0.034 |
| *Vegans* | 60 | 3.21 | 1.96 | 0.130 | – |
| *Lacto-ovo-vegetarians* | 53 | 3.92 | 2.27 | 0.094 | – |
| Omnivores | 59 | 0.11 | 0.07 | 0.973 | 0.973 |
| **Jaccard** | | | | | |
| Full cohort (pattern-adjusted) | 172 | 0.91 | 1.62 | 0.027 | – |
| Plant-based | 113 | 1.67 | 1.92 | 0.011 | 0.034 |
| *Vegans* | 60 | 2.46 | 1.49 | 0.064 | – |
| *Lacto-ovo-vegetarians* | 53 | 2.41 | 1.29 | 0.123 | – |
| Omnivores | 59 | 1.32 | 0.77 | 0.878 | 0.973 |
| **Bray-Curtis** | | | | | |
| Full cohort (pattern-adjusted) | 172 | 0.80 | 1.40 | 0.061 | – |
| Plant-based | 113 | 1.27 | 1.45 | 0.053 | 0.105 |
| *Vegans* | 60 | 1.89 | 1.13 | 0.260 | – |

| **Analysis** | **n** | **$R^2$ (%)** | **Pseudo-F** | **P** | **Q** |
|---|---|---|---|---|---|
| *Lacto-ovo-vegetarians* | 53 | 1.73 | 0.92 | 0.591 | – |
| Omnivores | 59 | 1.35 | 0.78 | 0.852 | 0.973 |

*Marginal PERMANOVA (adonis2, 9,999 permutations) of MPPF intake on microbiome composition, adjusted for sex, age and BMI. The full-cohort model additionally includes dietary pattern as a covariate, fitting a single slope across the three patterns; the stratified models are fitted independently within each stratum.*

*$R^2$ is the proportion of total variance and is therefore not comparable between the full cohort and a stratum; pseudo-F has an expectation of ~1 under the null regardless of sample size.*

*Q: Benjamini–Hochberg across the pre-specified family of 2 strata (plant-based, omnivores) × 4 metrics = 8 tests. Vegans and lacto-ovo-vegetarians (in italics) are reported as a sensitivity analysis and are not included in that family.*

*Aitchison, Euclidean distance on centred log-ratio-transformed abundances (primary metric); rclr, robust centred log-ratio; Jaccard, presence/absence; Bray-Curtis, relative abundance.*

Supplementary Table S5. Species of the ZOE health ranking and their association with MPPF intake, by dietary stratum.

| ZOE rank | Species | Family | Prev. PB (%) | OR PB (95% CI) | P PB | Q PB | Prev. OM (%) | OR OM (95% CI) | P OM |
|---|---|---|---|---|---|---|---|---|---|
| **Health-favourable species (50 species ranked most favourably)** | | | | | | | | | |
| 38 | *Vescimonas sp000435555* | Oscillospiraceae | 42 | 2.13 (1.33–3.43) | 0.002 | 0.080 | 46 | 1.30 (0.69–2.43) | 0.414 |
| 11 | *Eubacterium_F sp000433735* | Lachnospiraceae | 23 | 1.88 (1.17–3.01) | 0.009 | 0.133 | 31 | 0.97 (0.48–1.95) | 0.930 |
| 31 | *CAG-196 sp002102975* | Gastranaerophilaceae | 12 | 1.87 (1.08–3.24) | 0.025 | 0.133 | 25 | 1.28 (0.67–2.48) | 0.454 |
| 10 | *Suilimivivens sp000438155* | Lachnospiraceae | 33 | 1.82 (1.17–2.83) | 0.008 | 0.133 | 41 | 1.43 (0.77–2.65) | 0.262 |
| 19 | *CAG-115 sp003531585* | Ruminococcaceae | 30 | 1.69 (1.10–2.61) | 0.017 | 0.133 | 54 | 0.72 (0.39–1.33) | 0.298 |
| 20 | *Dysosmobacter segnis* | Oscillospiraceae | 67 | 1.69 (1.03–2.78) | 0.039 | 0.133 | 88 | 1.13 (0.46–2.76) | 0.790 |
| 30 | *CAG-170 sp000432135* | Oscillospiraceae | 50 | 1.69 (1.10–2.60) | 0.018 | 0.133 | 68 | 0.71 (0.38–1.33) | 0.278 |
| 35 | *Dysosmobacter sp900115635* | Oscillospiraceae | 23 | 1.64 (1.04–2.59) | 0.033 | 0.133 | 25 | 1.25 (0.65–2.38) | 0.505 |
| 8 | *UBA866 sp900543295* | Ruminococcaceae | 35 | 1.64 (1.06–2.53) | 0.025 | 0.133 | 46 | 0.78 (0.42–1.42) | 0.411 |
| 28 | *Hominifimenecus microfluidus* | Lachnospiraceae | 48 | 1.63 (1.05–2.51) | 0.028 | 0.133 | 44 | 1.26 (0.68–2.31) | 0.465 |
| 47 | *Scatosoma sp900542935* | Borkfalkiaceae | 18 | 1.60 (0.99–2.59) | 0.056 | 0.161 | 22 | 1.25 (0.63–2.50) | 0.526 |
| 5 | *CAG-170 sp900553545* | Oscillospiraceae | 52 | 1.59 (1.02–2.48) | 0.039 | 0.133 | 56 | 0.92 (0.52–1.63) | 0.770 |
| 16 | *QAKD01 sp003343965* | Anaerovoracaceae | 27 | 1.58 (1.03–2.42) | 0.036 | 0.133 | 36 | 0.83 (0.45–1.53) | 0.553 |
| 18 | *CAG-170 sp002404795* | Oscillospiraceae | 50 | 1.57 (1.02–2.42) | 0.041 | 0.133 | 58 | 1.06 (0.58–1.95) | 0.855 |
| 42 | *Faecousia sp000434635* | Oscillospiraceae | 48 | 1.56 (1.01–2.41) | 0.043 | 0.133 | 61 | 1.08 (0.60–1.97) | 0.792 |
| 2 | *CAG-269 sp003525075* | CAG-508 | 30 | 1.56 (1.03–2.38) | 0.037 | 0.133 | 37 | 1.81 (0.95–3.46) | 0.073 |
| 1 | *Hominicoprocola sp900317525* | Oscillospiraceae | 59 | 1.50 (0.97–2.32) | 0.068 | 0.168 | 71 | 0.85 (0.45–1.58) | 0.600 |
| 33 | *SFEL01 sp004557245* | CAG-138 | 41 | 1.50 (0.98–2.29) | 0.063 | 0.168 | 51 | 0.70 (0.39–1.29) | 0.256 |
| 39 | *JAGTTR01 sp018223385* | Lachnospiraceae | 58 | 1.47 (0.96–2.26) | 0.077 | 0.177 | 69 | 0.60 (0.31–1.14) | 0.119 |

| ZOE rank | Species | Family | Prev. PB (%) | OR PB (95% CI) | P PB | Q PB | Prev. OM (%) | OR OM (95% CI) | P OM |
|---|---|---|---|---|---|---|---|---|---|
| 27 | *Choladousia sp902363135* | Lachnospiraceae | 75 | 1.47 (0.88–2.45) | 0.138 | 0.235 | 81 | 0.96 (0.43–2.10) | 0.912 |
| 14 | *Suilimivivens sp. (SGB4893)* | Lachnospiraceae | 30 | 1.47 (0.95–2.26) | 0.081 | 0.177 | 20 | 1.14 (0.57–2.29) | 0.709 |
| 29 | *Faecousia sp958436045* | Oscillospiraceae | 37 | 1.45 (0.97–2.18) | 0.070 | 0.168 | 53 | 0.92 (0.51–1.66) | 0.793 |
| 4 | *Flemingibacterium sp900546315* | CAG-272 | 44 | 1.40 (0.94–2.09) | 0.100 | 0.209 | 54 | 0.81 (0.44–1.47) | 0.486 |
| 48 | *Faecivicinus sp900553265* | Aristaeellaceae | 26 | 1.39 (0.91–2.14) | 0.127 | 0.232 | 39 | 0.95 (0.52–1.73) | 0.870 |
| 7 | *Faecalibacterium prausnitzii_I* | Ruminococcaceae | 70 | 1.39 (0.87–2.23) | 0.166 | 0.263 | 81 | 1.05 (0.51–2.19) | 0.888 |
| 40 | *Lawsonibacter_A hominis* | Oscillospiraceae | 39 | 1.39 (0.93–2.08) | 0.105 | 0.211 | 39 | 0.94 (0.49–1.78) | 0.841 |
| 50 | *Woodwardibium sp900760275* | CAG-382 | 23 | 1.39 (0.86–2.25) | 0.182 | 0.279 | 27 | 0.86 (0.45–1.64) | 0.641 |
| 9 | *CAG-170 sp937967235* | Oscillospiraceae | 42 | 1.37 (0.92–2.04) | 0.117 | 0.225 | 61 | 0.87 (0.48–1.58) | 0.644 |
| 21 | *UBA11524 sp000437595* | Aristaeellaceae | 58 | 1.36 (0.89–2.08) | 0.160 | 0.263 | 63 | 0.76 (0.42–1.39) | 0.375 |
| 44 | *HGM12998 sp900756495* | Oscillospiraceae | 36 | 1.36 (0.91–2.02) | 0.131 | 0.232 | 37 | 1.26 (0.69–2.30) | 0.444 |
| 26 | *HGM14224 sp900761905* | HGM14224 | 29 | 1.34 (0.87–2.06) | 0.191 | 0.283 | 47 | 0.75 (0.40–1.39) | 0.356 |
| 37 | *CAG-269 sp001916005* | CAG-508 | 13 | 1.27 (0.75–2.16) | 0.381 | 0.515 | 20 | 1.51 (0.75–3.04) | 0.252 |
| 24 | *UBA3402 sp003478355* | Lachnospiraceae | 74 | 1.26 (0.77–2.04) | 0.358 | 0.499 | 71 | 1.46 (0.72–2.94) | 0.291 |
| 36 | *COE1 sp001916965* | Lachnospiraceae | 36 | 1.21 (0.81–1.79) | 0.352 | 0.499 | 29 | 1.24 (0.64–2.38) | 0.526 |
| 25 | *Eubacterium_F sp003491505* | Lachnospiraceae | 65 | 1.19 (0.77–1.81) | 0.434 | 0.571 | 81 | 0.90 (0.43–1.88) | 0.771 |
| 32 | *Lachnospira hominis* | Lachnospiraceae | 63 | 1.17 (0.78–1.76) | 0.456 | 0.583 | 61 | 1.39 (0.73–2.64) | 0.320 |
| 12 | *Roseburia_B zhanii* | Lachnospiraceae | 26 | 1.16 (0.75–1.80) | 0.510 | 0.635 | 31 | 1.43 (0.74–2.75) | 0.286 |
| 45 | *HGM11417 sp900761895* | HGM11417 | 13 | 1.15 (0.68–1.96) | 0.604 | 0.713 | 14 | 0.99 (0.40–2.45) | 0.978 |
| 43 | *Scatosoma sp900552625* | Borkfalkiaceae | 39 | 1.07 (0.72–1.58) | 0.746 | 0.837 | 49 | 1.14 (0.64–2.02) | 0.652 |
| 3 | *UBA11774 sp003507655* | Lachnospiraceae | 72 | 1.06 (0.68–1.65) | 0.803 | 0.859 | 80 | 1.27 (0.59–2.74) | 0.548 |

| ZOE rank | Species | Family | Prev. PB (%) | OR PB (95% CI) | P PB | Q PB | Prev. OM (%) | OR OM (95% CI) | P OM |
|---|---|---|---|---|---|---|---|---|---|
| 41 | *Ruminiclostridium_E siraeum* | Ruminococcaceae | 50 | 1.00 (0.68–1.46) | 0.988 | 0.988 | 61 | 1.23 (0.68–2.26) | 0.493 |
| 34 | *Roseburia_D yibonii* | Lachnospiraceae | 36 | 0.98 (0.66–1.47) | 0.940 | 0.961 | 31 | 1.14 (0.62–2.11) | 0.668 |
| 13 | *Instestinimonas_A celer* | Oscillospiraceae | 46 | 0.98 (0.67–1.45) | 0.937 | 0.961 | 36 | 1.71 (0.89–3.29) | 0.108 |
| 15 | *CAG-245 sp000435175* | CAG-508 | 33 | 0.95 (0.62–1.45) | 0.795 | 0.859 | 47 | 1.00 (0.56–1.79) | 0.990 |
| 22 | *Choladousia sp902363665* | Lachnospiraceae | 44 | 0.92 (0.62–1.38) | 0.697 | 0.801 | 49 | 1.11 (0.62–1.98) | 0.724 |
| 49 | *Alistipes communis* | Rikenellaceae | 81 | 0.88 (0.55–1.39) | 0.573 | 0.694 | 93 | – | – |
| 6 | *Clostridium lentum* | Clostridiaceae | 9 | – | – | – | 14 | 0.63 (0.25–1.58) | 0.323 |
| 17 | *V9D3004 sp900760345* | Lachnospiraceae | 6 | – | – | – | 5 | – | – |
| 23 | *UMGS621 sp900543915* | Borkfalkiaceae | 7 | – | – | – | 12 | 0.70 (0.26–1.86) | 0.478 |
| 46 | *CAJJNI01 sp905193085* | Lachnospiraceae | 8 | – | – | – | 14 | 1.58 (0.70–3.60) | 0.272 |
| **Health-unfavourable species (50 species ranked most unfavourably)** | | | | | | | | | |
| 27 | *Pseudoflavonifractor_A gallinarum* | Oscillospiraceae | 13 | 1.39 (0.80–2.42) | 0.246 | 0.810 | 20 | 1.42 (0.71–2.83) | 0.322 |
| 46 | *Eisenbergiella tayi* | Lachnospiraceae | 29 | 1.18 (0.77–1.80) | 0.442 | 0.883 | 19 | 1.02 (0.46–2.27) | 0.968 |
| 15 | *Eggerthella lenta* | Eggerthellaceae | 25 | 1.17 (0.73–1.87) | 0.503 | 0.891 | 32 | 1.12 (0.61–2.06) | 0.710 |
| 50 | *Pseudoscilispira sp020555605* | Oscillospiraceae | 15 | 1.12 (0.66–1.88) | 0.674 | 0.899 | 17 | 0.69 (0.30–1.57) | 0.371 |
| 17 | *Enterocloster aldenensis* | Lachnospiraceae | 20 | 1.11 (0.69–1.80) | 0.660 | 0.899 | 15 | 0.78 (0.34–1.82) | 0.569 |
| 24 | *Escherichia coli_F* | Enterobacteriaceae | 50 | 1.04 (0.70–1.55) | 0.838 | 0.921 | 44 | 1.07 (0.59–1.91) | 0.828 |
| 40 | *Enterocloster citroniae* | Lachnospiraceae | 42 | 1.02 (0.69–1.50) | 0.921 | 0.921 | 31 | 2.40 (1.16–4.95) | 0.018 |
| 14 | *Enterocloster bolteae* | Lachnospiraceae | 27 | 0.97 (0.63–1.50) | 0.897 | 0.921 | 22 | 1.23 (0.62–2.43) | 0.549 |
| 22 | *Enterocloster clostridioformis* | Lachnospiraceae | 18 | 0.97 (0.58–1.62) | 0.900 | 0.921 | 3 | – | – |
| 38 | *Thomasclavelia ramosa* | Coprobacillaceae | 12 | 0.95 (0.52–1.75) | 0.875 | 0.921 | 14 | 0.90 (0.37–2.16) | 0.809 |

| ZOE rank | Species | Family | Prev. PB (%) | OR PB (95% CI) | P PB | Q PB | Prev. OM (%) | OR OM (95% CI) | P OM |
|---|---|---|---|---|---|---|---|---|---|
| 10 | *Flavonifractor plautii* | Oscillospiraceae | 73 | 0.88 (0.57–1.37) | 0.577 | 0.891 | 75 | 1.13 (0.56–2.28) | 0.737 |
| 31 | *Hungatella effluvii* | Lachnospiraceae | 15 | 0.85 (0.47–1.53) | 0.579 | 0.891 | 3 | – | – |
| 5 | *Mediterraneibacter torques* | Lachnospiraceae | 41 | 0.83 (0.54–1.26) | 0.377 | 0.837 | 69 | 0.93 (0.50–1.73) | 0.824 |
| 39 | *CAG-274 sp000432155* | CAG-274 | 12 | 0.74 (0.39–1.43) | 0.373 | 0.837 | 7 | – | – |
| 12 | *Schaedlerella glycyrrhizinilytica_A* | Lachnospiraceae | 20 | 0.74 (0.43–1.28) | 0.283 | 0.810 | 32 | 1.79 (0.91–3.49) | 0.089 |
| 2 | *Enterocloster sp001517625* | Lachnospiraceae | 18 | 0.66 (0.36–1.19) | 0.168 | 0.671 | 7 | – | – |
| 21 | *Clostridium_AQ innocuum* | Erysipelotrichaceae | 21 | 0.58 (0.31–1.06) | 0.076 | 0.378 | 8 | – | – |
| 11 | *Ruminococcus_B gnavus* | Lachnospiraceae | 37 | 0.54 (0.32–0.91) | 0.021 | 0.210 | 39 | 1.17 (0.64–2.14) | 0.605 |
| 43 | *Blautia_A fusiformis* | Lachnospiraceae | 76 | 0.53 (0.34–0.84) | 0.007 | 0.133 | 88 | 0.34 (0.10–1.14) | 0.080 |
| 35 | *Clostridium_AP scindens* | Lachnospiraceae | 12 | 0.45 (0.19–1.08) | 0.073 | 0.378 | 10 | 1.81 (0.73–4.49) | 0.204 |
| 1 | *Faecalimonas umbilicata* | Lachnospiraceae | 3 | – | – | – | 3 | – | – |
| 3 | *Blautia hansenii* | Lachnospiraceae | 0 | – | – | – | 2 | – | – |
| 4 | *Amedibacillus dolichus* | Erysipelotrichaceae | 1 | – | – | – | 3 | – | – |
| 6 | *Blautia_A sp900547615* | Lachnospiraceae | 0 | – | – | – | 0 | – | – |
| 7 | *Blautia sp900541955* | Lachnospiraceae | 4 | – | – | – | 0 | – | – |
| 8 | *Blautia_A caecimuris* | Lachnospiraceae | 2 | – | – | – | 5 | – | – |
| 9 | *Brotaphodocola sp. (SGB4746)* | Lachnospiraceae | 7 | – | – | – | 19 | 1.09 (0.53–2.26) | 0.807 |
| 13 | *Otoolea sp000435655* | Lachnospiraceae | 0 | – | – | – | 12 | 0.86 (0.34–2.19) | 0.755 |
| 16 | *Blautia argi* | Lachnospiraceae | 1 | – | – | – | 7 | – | – |
| 18 | *Ruminococcaceae sp. (SGB79883)* | Ruminococcaceae | 4 | – | – | – | 0 | – | – |
| 19 | *Blautia_A wexlerae_B* | Lachnospiraceae | 96 | – | – | – | 100 | – | – |

| ZOE rank | Species | Family | Prev. PB (%) | OR PB (95% CI) | P PB | Q PB | Prev. OM (%) | OR OM (95% CI) | P OM |
|---|---|---|---|---|---|---|---|---|---|
| 20 | *Anaerostipes caccae* | Lachnospiraceae | 0 | – | – | – | 0 | – | – |
| 23 | *JANFXS01 sp024460105* | Anaerotignaceae | 8 | – | – | – | 3 | – | – |
| 25 | *Egerieimonas sp900552095* | Lachnospiraceae | 0 | – | – | – | 2 | – | – |
| 26 | *Blautia sp000432195* | Lachnospiraceae | 0 | – | – | – | 3 | – | – |
| 28 | *Lawsonibacter sp902363045* | Oscillospiraceae | 4 | – | – | – | 5 | – | – |
| 29 | *(SGB10115)* | – | 4 | – | – | – | 5 | – | – |
| 30 | *Dorea_B phocaeensis* | Lachnospiraceae | 0 | – | – | – | 2 | – | – |
| 32 | *Faecalimonas sp029983955* | Lachnospiraceae | 7 | – | – | – | 5 | – | – |
| 33 | *Sellimonas intestinalis* | Lachnospiraceae | 7 | – | – | – | 5 | – | – |
| 34 | *(SGB8163)* | – | 0 | – | – | – | 0 | – | – |
| 36 | *Phocaeicola vulgatus* | Bacteroidaceae | 93 | – | – | – | 86 | 0.76 (0.33–1.75) | 0.519 |
| 37 | *Bacteroides uniformis* | Bacteroidaceae | 96 | – | – | – | 97 | – | – |
| 41 | *Lactococcus cremoris* | Streptococcaceae | 3 | – | – | – | 7 | – | – |
| 42 | *Longicatena caecimuris* | Erysipelotrichaceae | 7 | – | – | – | 10 | 0.49 (0.15–1.64) | 0.248 |
| 44 | *Blautia celeris* | Lachnospiraceae | 4 | – | – | – | 3 | – | – |
| 45 | *Metalachnospira sp900548595* | Anaerotignaceae | 10 | – | – | – | 10 | 0.38 (0.11–1.24) | 0.108 |
| 47 | *Fimivicinus sp900544375* | Acutalibacteraceae | 7 | – | – | – | 5 | – | – |
| 48 | *Dorea_D sp000433535* | Lachnospiraceae | 0 | – | – | – | 10 | 0.69 (0.24–1.98) | 0.491 |
| 49 | *Streptococcus anginosus* | Streptococcaceae | 2 | – | – | – | 2 | – | – |

*Upper block: the 50 species ranked most favourably associated with cardiometabolic health markers in the ZOE Microbiome Health Ranking 2025 (Asnicar et al. 2026); lower block: the 50 ranked most unfavourably. Within each block species are ordered by the odds ratio in plant-based participants.*

*OR: odds ratio for species presence per SD of MPPF intake, from logistic regression adjusted for sex, age, BMI and sequencing depth, fitted separately in plant-based (PB, n = 113) and omnivore (OM, n = 59) participants. Species with prevalence <10% or >90% within a stratum were not tested (–).*

*Q: Benjamini-Hochberg within each set and stratum. Species-level results are exploratory; the formal test is the set-level analysis.*
*Taxonomy: GTDB release 220 names from the MetaPhlAn 4 SGB-to-GTDB mapping (mpa_vJan25_CHOCOPhlAnSGB_202503); species without a GTDB species name are shown as genus (or family) and SGB identifier.*

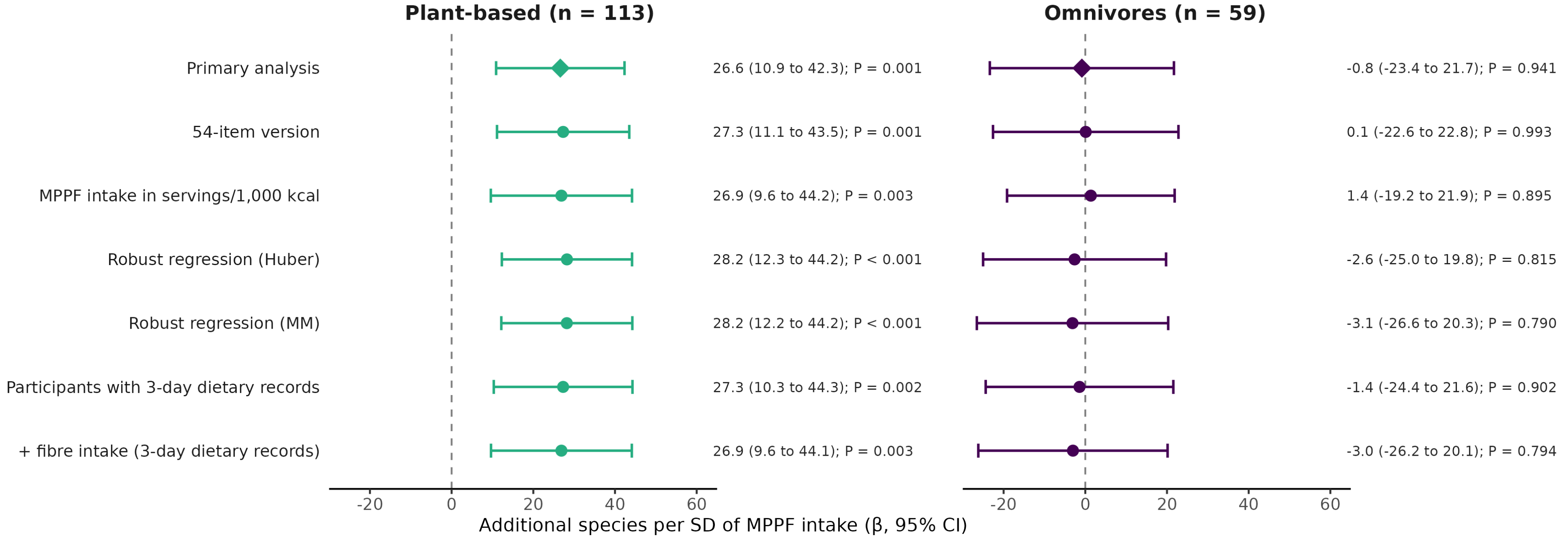


**Supplementary Fig. S1.** Sensitivity analyses of the association between MPPF intake and observed species richness. Points show β (additional species per SD of the corresponding exposure) with 95% confidence intervals, from models adjusted for sex, age, BMI and sequencing depth, fitted separately in plant-based participants (lacto-ovo-vegetarians and vegans combined; n = 113) and omnivores (n = 59); values are given on the right. Diamonds indicate the primary analysis (SD = 209 g/1,000 kcal). Sensitivity analyses used the 54-item version of MPPF intake, MPPF intake expressed in servings/1,000 kcal, and robust regression (Huber M-estimation and MM-estimation). The 54-item and servings-based versions were standardised to their own SD; all other analyses used the primary measure. Analyses in participants with 3-day dietary records were restricted to that subsample (100 plant-based participants and 58 omnivores) and were fitted without and with additional adjustment for total fibre intake (g/1,000 kcal) estimated from the records. Full results are given in Supplementary Table S3.

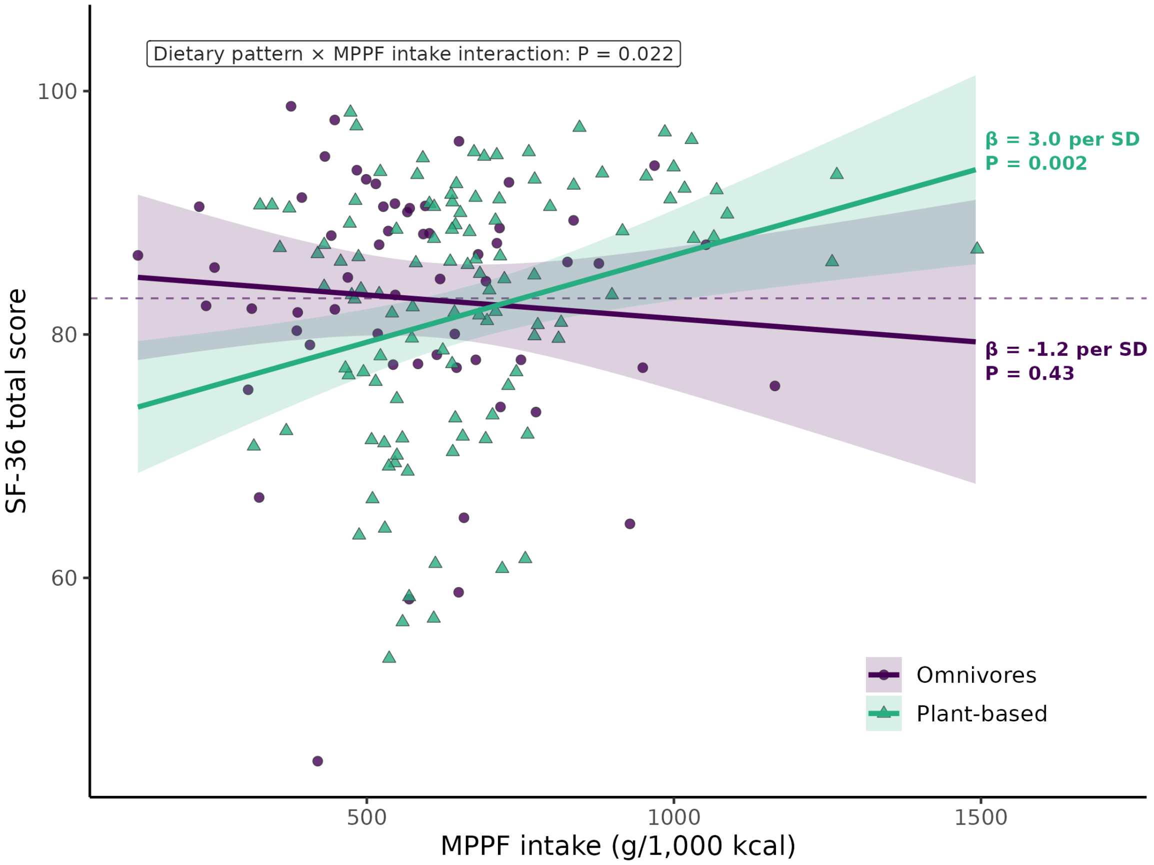


**Supplementary Fig. S2.** MPPF intake and SF-36 total score by dietary pattern. Lines and shaded bands show predicted values and 95% confidence intervals from a linear model including a dietary pattern × MPPF intake interaction, adjusted for sex, age and BMI; points are individual participants (circles, omnivores, n = 59; triangles, plant-based participants, n = 113). β values are within-group slopes per SD of MPPF intake (209 g/1,000 kcal) from stratified models with the same covariates. The dashed line marks mean SF-36 total score in omnivores. Higher scores indicate better health-related quality of life.
